# Release site plasticity via Unc13A regulatory domains mediates synaptic short-term facilitation and homeostatic potentiation

**DOI:** 10.1101/2022.11.16.516613

**Authors:** Meida Jusyte, Natalie Blaum, Mathias A. Böhme, Manon M. M. Berns, Alix E. Bonard, Janus R. L. Kobbersmed, Alexander M. Walter

## Abstract

Chemical synaptic transmission relies on neurotransmitter release from presynaptic release sites and on transmitter-sensing by the postsynaptic cell. Presynaptic plasticity increasing neurotransmitter release achieves two fundamental nervous system functions: It tunes some synapses to be more responsive to millisecond repetitive activation and it maintains signals when postsynaptic transmitter sensitivity is reduced. How enhanced neurotransmitter release is achieved in these phenomena, termed short-term facilitation and homeostatic potentiation, remains unknown. We combine mathematical modeling and experimental analysis of *Drosophila* neuromuscular junction model synapses to elucidate the molecular mechanisms underlying these forms of plasticity. Our results indicate that both phenomena depend on a rapid increase in the participation of neurotransmitter release sites which is controlled by the regulatory domains of the evolutionarily conserved (M)Unc13A protein that bind Ca^2+^/Calmodulin and diacylglycerol. Mutation of the Calmodulin binding (CaM) domain increased baseline transmission and impaired both short-term facilitation and acute homeostatic potentiation. Mathematical modeling indicated that these defects result from too many release sites participating at rest combined with the inability to plastically further increase their number. Super-resolution microscopy revealed that this coincided with a redistribution of Unc13A’s functionally essential MUN domain closer to the synaptic plasma membrane, which may constitute the molecular switch to increase release site participation. Similar consequences (enhanced baseline transmission, block of both short-term facilitation and homeostatic potentiation) were caused by the acute pharmacological activation of the C1 domain of wildtype Unc13A using phorbol esters. This treatment had no effect on Unc13A CaM domain mutants, indicating that both the CaM and C1 domains activate a binary release site switch. Thus, our findings indicate that Unc13A regulatory domains are tuned to integrate a multitude of signals on various timescales to switch release site participation for synaptic plasticity.

## Introduction

Chemical synaptic transmission depends on action potential (AP) induced Ca^2+^ influx triggering the fusion of synaptic vesicles (SVs) with the plasma membrane at release sites of presynaptic active zones (AZs). The released Neurotransmitters (NTs) are subsequently detected by postsynaptic receptors (Sudhof, 2013). Both the pre- and post-synaptic sides can be involved in synaptic plasticity, the modulation of transmission strength (Citri and Malenka, 2008). Short-term facilitation (STF), the transient increase in synaptic responses upon repetitive stimulation on the millisecond timescale, primarily relies on presynaptic mechanisms and forms a basis of the nervous system for temporal information processing (Abbott and Regehr, 2004; Fioravante and Regehr, 2011). Changes in the vesicular release probability (pV_r_) to AP stimulation may contribute to STF (Zucker and Regehr, 2002), but if pV_r_s are heterogeneous (e.g. which is likely often the case as distances between SVs and Ca^2+^ channels vary), changes in pV_r_ alone are insufficient to achieve the strong STF experimentally observed (Kobbersmed et al., 2020). More recently, changes of release site/docking site occupation have been proposed to mediate STF (Kobbersmed *et al*., 2020; Kusick et al., 2020; Lin et al., 2022; Neher and Taschenberger, 2021; Pulido and Marty, 2018; Silva et al., 2021), but molecular details are still lacking.

Longer-term synaptic plasticity is essential to stabilize information flow and form memories (Citri and Malenka, 2008; Nicoll and Roche, 2013). Presynaptic homeostatic potentiation (PHP) is an evolutionarily conserved mechanism to enhance AP-induced NT release to ensure transmission in cases postsynaptic NT sensitivity is reduced (Davis and Muller, 2015). PHP is acutely induced (and observed after minutes) by the pharmacological inhibition of NT receptors at central mammalian synapses and the *Drosophila melanogaster* neuromuscular junction (NMJ) (Delvendahl et al., 2019; Frank et al., 2006). Chronic mutation of glutamate receptors of the NMJ similarly reduces NT sensitivity which is offset by a sustained (life-long) PHP (Frank *et al*., 2006). While acute and chronic PHP similarly enhance NT release, the underlying mechanisms differ: Longer-term PHP depends on cellular transport reactions and coincides with the local enrichment of synaptic components (Böhme et al., 2019; Goel et al., 2019; Weyhersmuller et al., 2011). In contrast, acute PHP on the few minutes’ timescale is achieved independently of cellular transport, indicating that rapid PHP relies on the available synaptic material which is “switched” to a state more permissive for NT release (Böhme *et al*., 2019). Yet how this is molecularly achieved is unknown.

The evolutionarily conserved (M)Unc13 proteins are essential for NT release and mediate the physical attachment of SVs to the plasma membrane (SV docking) and their molecular maturation to become responsive to the Ca^2+^ stimulus (SV priming) (Augustin et al., 1999; Betz et al., 1998; Bohme et al., 2016; Imig et al., 2014; Siksou et al., 2009; Varoqueaux et al., 2002). Unc13 proteins localize in defined clusters to generate the SV release sites, a property which is conserved across species (*Drosophila*/mouse/*C. elegans*) (Hu et al., 2013; Reddy-Alla et al., 2017; Sakamoto et al., 2018; Zhou et al., 2013). The (M)Unc13 proteins contain several evolutionarily conserved domains which have been shown to influence synaptic transmission (Dittman, 2019; Rizo and Rosenmund, 2008). Specifically, mutations of the Ca^2+^/Calmodulin interaction domain (CaM domain) or the Ca^2+^/phosphoinositide-binding C2B domain of murine Munc13-1 alter short-term plasticity (Junge et al., 2004; Lipstein et al., 2021; Reddy-Alla *et al*., 2017; Shin et al., 2010). Moreover, mutation of the protein’s C1 domain or its pharmacological activation with the diacylglycerol (DAG) analog phorbol ester (PMA) enhance neurotransmitter release (Basu et al., 2007; Betz *et al*., 1998; Maruyama and Brenner, 1991; Rhee et al., 2002; Schotten et al., 2015). However, how these manipulations exactly change transmitter release and under which biological circumstances this is relevant, remains unclear.

Here we investigate the relevance of *Drosophila* Unc13A regulatory domains in release site-based plasticity on the timescales of milliseconds and minutes. Mathematical modeling was used to predict the effects of impaired release site plasticity to fast, repetitive (paired-pulse) AP-stimulation. The model predicted that effects on synaptic responses should be best visible when varying extracellular Ca^2+^ concentrations. Electrophysiological recordings of the *Drosophila* larval NMJ revealed that mutation of the Unc13A CaM domain resulted in increased baseline transmission and loss of STF at low Ca^2+^, a phenotype consistent with constantly high release site participation and a consequent loss of plastic release site activation. Super-resolution STED microscopy was used to investigate whether this coincided with a redistribution of synaptic Unc13A. This identified a small shift of the catalytically active MUN domain towards the synaptic plasma membrane, indicating that a confirmational change may underly Unc13 and hence release site switching. A relevance on the minutes’ timescale was investigated by acutely challenging Unc13A CaM mutant synapses with pharmacological inhibition of NT receptors which demonstrated that the normally observed PHP upon this treatment was lost, consistent with release-site based plasticity underlying both STF and PHP. Functional synergism between Unc13A regulatory domains was investigated by pharmacologically targeting the Unc13A C1 domain using phorbol esters. This revealed similar phenotypes as mutation of the CaM domain (increased baseline transmission and abolished STF at low extracellular Ca^2+^, block of PHP) and no further enhancement was seen when combining both manipulations, indicating activation of either domain induces the same downstream release site activation. Thus, acute regulation of release site participation is a powerful presynaptic plasticity mechanism for short-term plasticity and immediate homeostasis and Unc13A regulatory domains can regulate this by integrating a variety of intracellular signals.

## Results

### Rapid regulation of neurotransmitter release sites as a mechanism of short-term facilitation

STF might rely on activity-dependent increases in the number of participating release sites between stimuli (Pulido and Marty, 2018). We recently proposed a facilitation model where the occupation of release sites increases upon AP stimulation by Ca^2+^-dependent stabilization of the primed vesicle state (Kobbersmed *et al*., 2020)(simplified model scheme in Fig 1A). Our model relies on the assumption that SV priming is fast but reversible and that SV unpriming is slowed by Ca^2+^. In facilitating synapses, unpriming commences similarly fast (or faster) as forward priming, resulting in incomplete release site occupation which limits NT release with the first AP stimulus. However, following the AP, Ca^2+^ accumulates in the synapse and slows unpriming (Kobbersmed *et al*., 2020). This increases release site occupation (Fig 1B_1_, B_2_) allowing more NT to be released with a second AP (Fig. 1B_3_), resulting in STF. The strongest facilitation is achieved when initial release site occupation is low, and many additional sites are populated between stimuli. This is the case when intracellular Ca^2+^ is low - and unpriming fast-at rest (Fig. 1A,B_1_,B_2_,B_3_). Because elevating the extracellular Ca^2+^ concentration likely also elevates its intracellular levels, this model predicts that the largest STF will be observed when extracellular Ca^2+^ levels are low, while increasing external Ca^2+^ will diminish facilitation by this mechanism (Kobbersmed *et al*., 2020). At high extracellular Ca^2+^ concentrations the model predicts that most release sites will be occupied already at rest (Fig. 1C_1_,C_2_). Furthermore, the release probability at occupied release sites is increased in this condition (APs cause more Ca^2+^ influx). Together, these two mechanisms lead to more NT release at the first AP (Fig. 1C_1_,C_3_). If a second AP triggers NT release before all of the emptied release sites are repopulated, less NT will be released, resulting in short-term depression (STD) (Fig. 1C_1_,C_3_). Synaptic transmission is typically assessed by recording from the postsynaptic cell which is why we used our model to predict AP-evoked Excitatory Postsynaptic Currents (eEPSCs) using the typical postsynaptic effect NT release from a single SV has and the rate with which SVs are released (Fig. 1D). Quantification of the peak eEPSC values revealed their increase and later saturation as external Ca^2+^ levels rose (Fig. 1E). An experimental quantitative measure of STF is the paired-pulse-ratio (PPR), the ratio of the amplitude of the second eEPSC response divided by the first which is greater than one for STF and smaller than one for STD. Our model predicts a gradual shift from STF to STD with increasing extracellular Ca^2+^ (Fig. 1E-F).

**Fig. 1:**
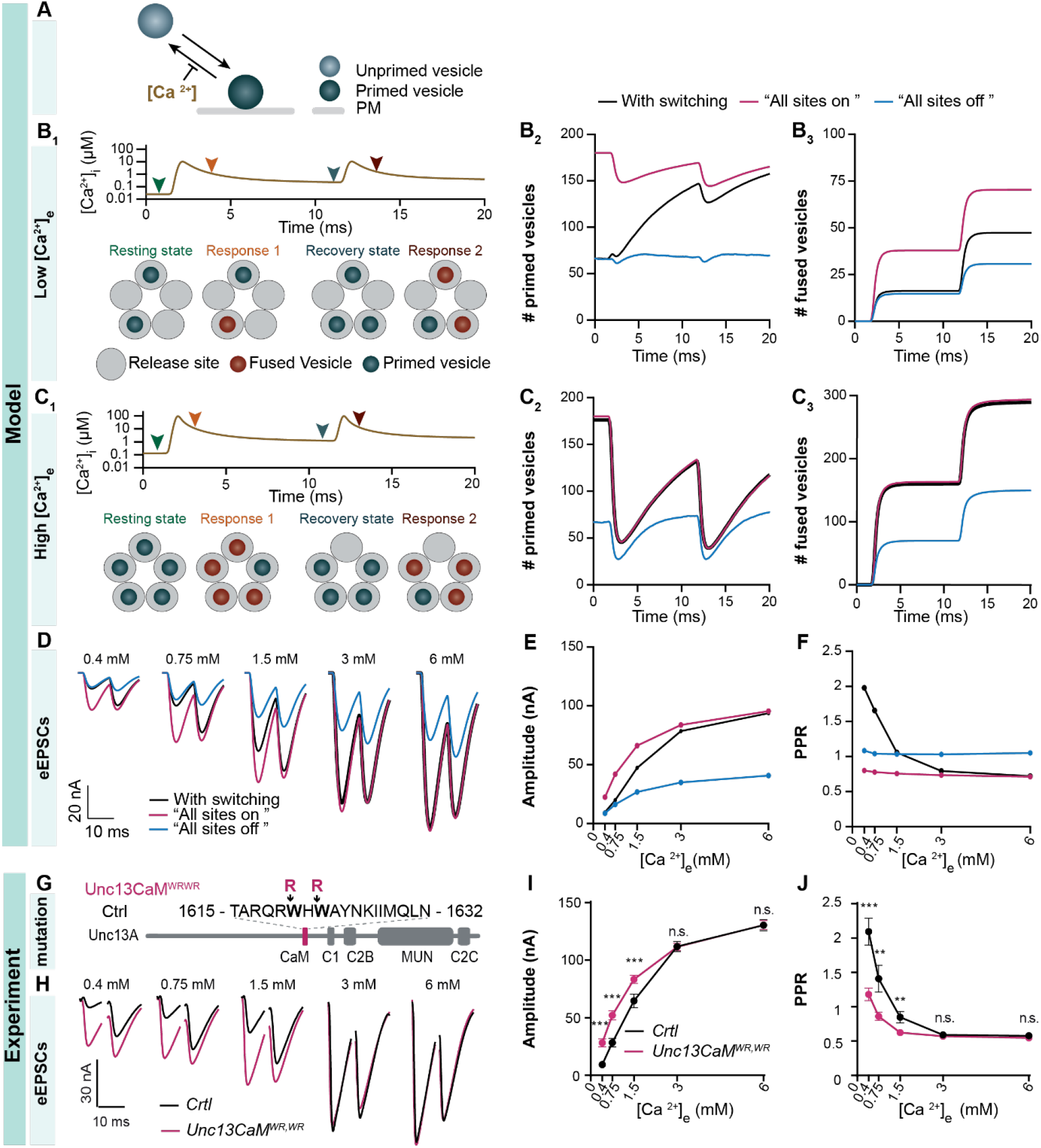
Constantly high release site occupation causes enhanced synaptic transmission and STD at low and intermediate Ca^2+^ concentrations, which can be observed in the Unc13 CaM^wr,wr^ mutant. (**A**) Simplified cartoon of a model in which vesicular unpriming is reduced with increasing internal Ca^2+^ concentrations as in Kobbersmed et al., 2020. (**B**_**1**_) (Top) Ca^2+^ transient (induced by an AP) at 120 nm distance from a Ca^2+^ source corresponding to a paired pulse stimulation (10 ms interstimulus interval) at a low extracellular Ca^2+^ concentration (0.4 mM). Arrows indicate time points that are illustrated in the bottom panel. (Bottom) Representative cartoon of five hypothetical release sites (gray circles) containing primed (blue circles) and fused (orange circles) vesicles at different time points during a paired pulse stimulation. The resting state (green arrow) corresponds to the state of release sites before the first AP. Response 1 (orange arrow) represents the release sites configuration just after the first AP. The second recovery state (blue arrow) and response 2 (red arrow) refer to the state of the release site just before and after the second AP, respectively. (**B**_**2**_) Mean number of primed vesicles across all 180 release sites at low extracellular Ca^2+^ concentration (0.4 mM) of the model over time during a paired pulse stimulation, for a model with Ca^2+^-dependent unpriming (with switching condition, black curve), and two different conditions in which unpriming is Ca^2+^-independent. The “All sites *on*” condition (pink curve) has no unpriming (r=0) and the “All sites *off*” condition (blue curve) has constitutively fast unpriming (r=1) (seeFig. S1). (**B**_**3**_) Cumulative number (mean) of fused vesicles over time for the condition with release site switching (black curve), “All sites *on*” (pink curve) and “All sites *off*” (blue curve) conditions, for low extracellular concentration (0.4 mM). (**C**) Same as B but for high extracellular Ca^2+^ concentration of 6 mM. (**D**) Simulated average eEPSCs for different extracellular Ca^2+^concentrations simulated for a model with release site switching (black trace), and with the “All sites *on*” (pink trace) and “All sites *off”* conditions (blue trace). (**E**) Mean amplitude of the first eEPSC for the release site switching (black), “All sites *on*” (pink), “All sites *off*” (blue) conditions. (**F**) Paired-pulse ratio computed from simulated eEPSCs for the release site switching (black), “All sites *on*” (pink), “All sites *off*” (blue) conditions, for different extracellular Ca^2+^ concentrations. Mean model predictions are computed over 200 stochastic model repetitions. **(G)** Scheme of the Unc13A protein indicating the regulatory Ca^2+^/Calmodulin binding CaM-domain, the DAG/PMA-binding C1 domain, the Ca^2+^/phosphoinosite-binding C2B domain, the functionally essential MUN domain and the C2C domain implicated in SV association. and the Ca, C2B, DAG/PMA binging C1 domain. Note that unlike in mammals and C. elegans, the N-terminal C2A domain is missing in flies. The amino acid change in the here analyzed CaM^W1620R,W1622R^ mutant is shown. (**H**) Representative example traces of evoked responses to paired APs (10 ms interstimulus interval) of a single *Ctrl* (black) and *CaM*^*W1620R,W1622R*^ (magenta) cells recorded at increasing Ca^2+^ concentrations (from 0.4 to 6 mM). (**I+J**) Quantification of first eEPSCs (I) and PPRs (J) in *Ctrl* (black) and *CaM*^*W1620R,W1622R*^ mutant (magenta). Genotypes (see methods for details): *Ctrl*: Unc13A and -B null animals expressing wildtype Unc13A; *CaM*^*W1620R,W1622R*^ mutant: Unc13A and -B null animals expressing the Unc13A *CaM*^WRWR^ mutant. Data of *Ctrl* (black) and *CaM*^*W1620R,W1622R*^ mutant are averaged across animals and shown with ± SEM. Number of animals (N): N(*Ctrl*)=20, N(*CaM*^*W1620R,W1622R*^ mutant)=30. Data depicts mean values ± s.e.m. Statistical analysis with multiple unpaired parametric t-tests. Asterisks indicate * p ≤ 0.05; ** p ≤ 0.01; *** ≤ 0.001.

What might be the consequences of losing this release-site based short-term plasticity? We evaluated this by simulating hypothetical conditions that abolish the Ca^2+^-dependent changes of the unpriming rate which may cause unpriming to be constitutively slow or fast (Fig. S1). Constitutively fast unpriming results in constantly low site occupation and diminished synaptic responses (Fig. 1B_2_-F). The effect becomes particularly clear under conditions of high extracellular Ca^2+^ where the release site occupation increases in the normal condition due to a reduced unpriming rate (Fig. 1A,C_2_-F). In contrast, constitutively slow unpriming leads to unnaturally high site occupation at low extracellular Ca^2+^ where it, compared to the control condition, causes enhanced transmission while responses were similar at high extracellular Ca^2+^ where unpriming is slowed in the control condition (Fig. 1B_2_-F). In both cases, the normally observed STF at low Ca^2+^ is abolished, making the PPR profile largely independent of the external Ca^2+^ concentration (Fig. 1F). Thus, while both hypothetical conditions similarly affect STP behavior, differences are observed on initial synaptic responses when probed at high or low extracellular Ca^2+^ (Fig. 1E,F).

### Mutation of the Unc13A Calmodulin binding domain enhances synaptic transmission and abolishes STF at low extracellular Ca^2+^

We sought to investigate potential molecular mechanisms of the activity-dependent enhancement of release site occupation. Due to its pivotal role in release site function, its conserved regulatory domains and its function in priming/unpriming, Unc13A is a prime target for this plasticity (Camacho et al., 2017; He et al., 2017; Lipstein *et al*., 2021; Liu et al., 2016; Michelassi et al., 2017; Reddy-Alla *et al*., 2017). Mutation of the murine Munc13-1 Calmodulin binding domain increased STD in hippocampal neurons (Junge *et al*., 2004) and we wondered to which extent this might be due to constitutively high-or low release site occupation. We therefore compared eEPSCs measured by two electrode voltage clamp (TEVC) recordings at the *Drosophila* 3^rd^ instar larval neuromuscular junction (NMJ) in control flies expressing the Unc13A wildtype protein to ones bearing a corresponding mutation of the Unc13A CaM domain (see methods for exact genotypes) (Reddy-Alla *et al*., 2017). Confocal analysis confirmed similar expression and targeting of mutant and wildtype proteins to NMJ AZs (Fig. S2). Motivated by our modelling approach, we recorded synaptic responses to paired AP stimulation (10 ms apart) while varying the extracellular Ca^2+^ concentration (Fig. 1G, H). This revealed much larger responses of the CaM mutant compared to controls at low (0.4 mM, 0.75 mM, 1.5 mM)-but similar responses at high (3 mM, 6 mM) extracellular Ca^2+^ concentration (Fig. 1I). Moreover, the typical STF at low Ca^2+^ observed in animals expressing wildtype Unc13A, was lost in the mutants (Fig. 1H, J). Because of the strong resemblance of these characteristics with the model predictions for constitutively slow unpriming (compare Fig. 1E-F and I-J), these data are consistent with the CaM mutation resulting in constantly high release site occupation which bypasses the need for Ca^2+^ elevation. Thus, millisecond changes of release site occupation constitute a powerful STF mechanism and Unc13A domains might regulate this.

### Mutation of Unc13A’s CaM domain results in closer membrane apposition of its MUN domain

Above data are consistent with the mutation of the CaM domain resulting in higher release site occupation. What might be the molecular mechanism underlying this? (M)Unc13 proteins have an extended structure and conformational changes were proposed to change their function (Camacho et al., 2021; Grushin et al., 2022). The conserved C-terminal MUN domain is essential and sufficient to promote SV priming (Basu et al., 2005; Stevens et al., 2005) and therefore conformational changes that affect its relative orientation to the plasma membrane could well contribute to changes in release site occupation. We therefore investigated to which extent this domain was repositioned upon mutation of the CaM domain using super-resolution STED microscopy. AZs of 3^rd^ instar larvae were stained using antibodies recognizing the Unc13A MUN domain (Reddy-Alla *et al*., 2017) and the C-terminus of the ELKS-family scaffolding protein Bruchpilot (BRP) (Kittel et al., 2006). The BRP epitope detected here is located at the roof of the pedestal with maximal distance from the AZ plasma membrane (Fig. 2A) (Fouquet et al., 2009). STED imaging revealed the typical ring-like structures for AZs in top view (yellow boxes in Fig. 2A-E) (Kittel *et al*., 2006). Analysis of top view AZ images can be used to investigate possible changes in the distance to the AZ center, where voltage gated Ca^2+^ channels are located (Bohme *et al*., 2016; Fouquet *et al*., 2009). To quantitatively investigate this, we centered many individual AZs based on their BRP signal (see methods) and averaged those to compare potential differences in the distribution of BRP and Unc13A in both genotypes (Fig. S3). The analysis confirmed the overall ring-like distribution of the BRP C-terminus and a similar, but more AZ-central distribution of Unc13A MUN domain (Fig. 2A-E, Fig. S3). No differences between the two genotypes were observed (Fig. S3), indicating that mutation of the CaM domain does not grossly affect AZ morphology or the distance between Unc13A release sites and the AZ centrally positioned voltage gated Ca^2+^ channels.

**Fig. 2:**
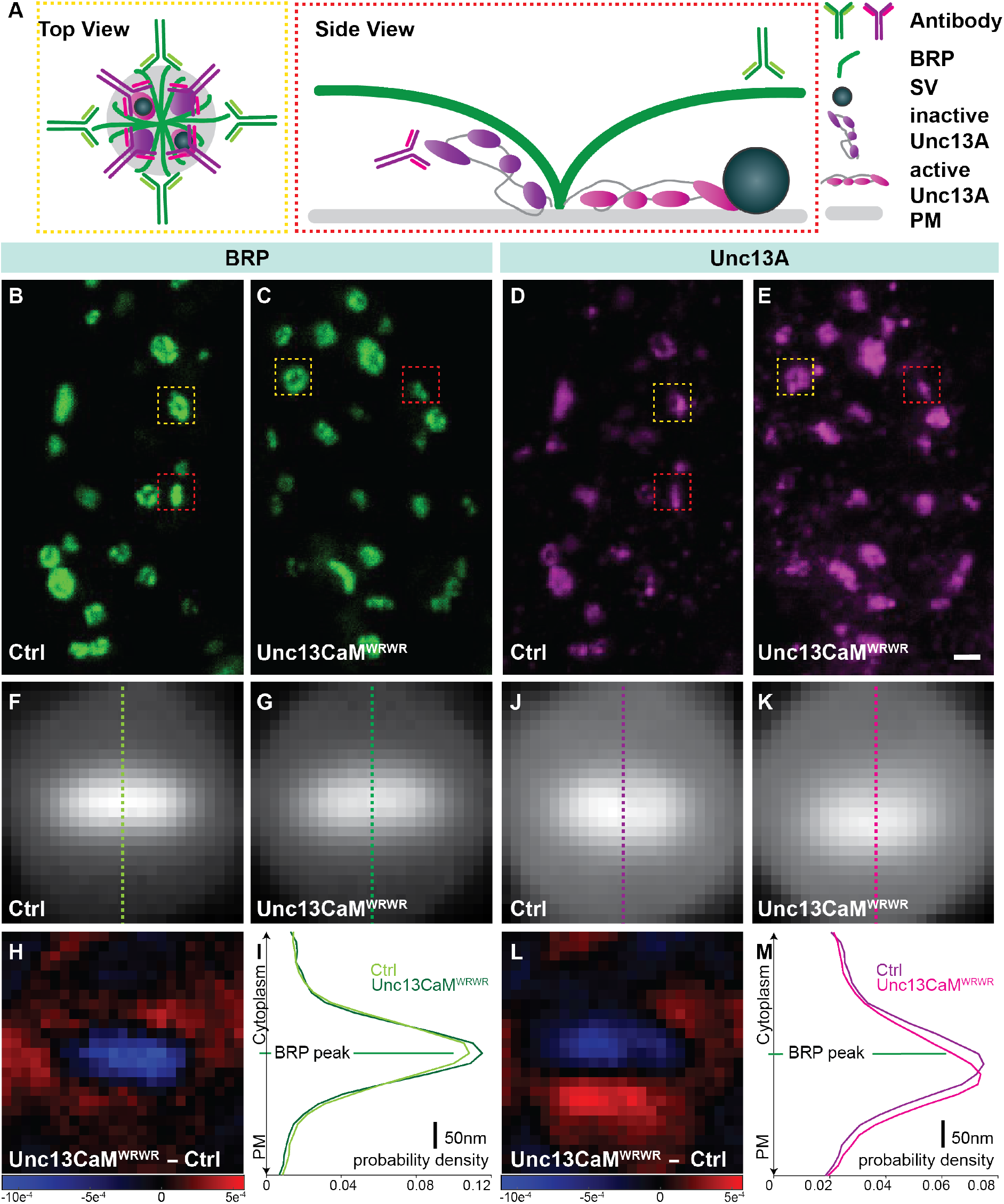
Unc13A CaM mutation positions the Unc13A MUN domain closer to the plasma membrane. **(A)** Schematics of top (yellow dashed box) and laterally (red dashed box) viewed AZ with BRP in green and Unc13A in purple (inactive) or pink (active). Schematic antibodies indicating binding epitopes (green: NC82/Bruchpilot; purple: MUN-Domain). **(B-E)** STED images of muscle 4 NMJs of segment A2–4 from third-instar larvae of the displayed genotypes labeled with antibodies recognizing BRP (green) and Unc13A purple). Red dashed boxes show laterally viewed AZ. Yellow dashed boxes show top-viewed AZ. **(F-G)** Average, and normalized (total sum of the 25×25 pixels equal to 1), BRP signal from single AZ images of the displayed genotypes obtained after aligning the BRP signal of all individual ROIs to the center (see methods). Ctrl and Unc13CaM^wrwr^ images are displayed on the same scale. Dashed vertical lines through the midpoint of the image show which intensity values were used to generate line profiles (I). Black edges are caused by rotation of the ROIs during the alignment procedure. **(H)** Subtracted image of the averaged and normalized BRP signal in the Unc13A CaM^WRWR^ mutant minus the obtained from control animals. **(J-K)** Average, and normalized (total sum of the 25×25 pixels equal to 1), Unc13 signal of the displayed genotypes obtained after aligning the BRP signal of all individual ROIs to the center, the Unc13A channel “followed” the alignment of BRP (see methods). Ctrl and Unc13CaM^wrwr^ images are displayed on the same scale. Dashed vertical lines through the midpoint of the image show which intensity values were used to generate line profiles (M). **(L)** Subtracted image of the averaged and normalized Unc13A signal in the Unc13A CaM^WRWR^ mutant minus the obtained from control animals. The image is displayed at the same scale as in H. **(I-M)** Line profiles of BRP and Unc13A for the displayed genotypes. Profiles are scaled such that area under the curve equals 1. Genotypes (see methods for details): *Ctrl*: Unc13A and -B null animals expressing wildtype Unc13A; *CaM*^*W1620R,W1622R*^ mutant: Unc13A and -B null animals expressing the Unc13A *CaM*^WRWR^ mutant. Number of analyzed AZs (*ν*), NMJs(n) and animals (N): *ν*(*Unc13CaM*^*WRWR*^)=63, n(*Unc13CaM*^*WRWR*^)=7, N(*Unc13CaM*^*WRWR*^)= 4; *ν*(Ctrl)=67, n(Ctrl)=9, N(Ctrl)=4. Scale bar: 400 nm. Size of individual, averaged and subtracted AZ images: 25×25 pixel, pixel size=20 nm.

To investigate whether mutation of the CaM domain affected the distance of the Unc13A MUN domain to the plasma membrane, we analyzed AZs observed in side-view (Fig. 2A-D) (Fouquet *et al*., 2009; Reddy-Alla *et al*., 2017). A quantitative comparison between genotypes was performed by using the BRP signal to center and rotating many individual AZ images such that they ran from cytosol to plasma membrane from top to bottom (see methods). The same translations and rotations were applied to the Unc13A channels before all images were averaged and compared. This revealed that the mean fluorescence distribution profile from cytosol towards the plasma membrane was highly similar for BRP in both genotypes (Fig. 2F,G,I). Also subtracting the average images obtained in either genotype from one another revealed no major change in fluorescence distribution, indicating that mutation of the Unc13A CaM domain did not grossly affect synaptic BRP distribution (Fig. 2H). Consistent with a previous analysis of the wildtype protein (Reddy-Alla *et al*., 2017), the fluorescence in the Unc13A channel peaked closer to the plasma membrane in both genotypes (compare Fig2I to 2M). Furthermore, this analysis revealed that the area of high fluorescence was shifted towards the plasma membrane upon mutation of the Unc13A CaM domain (Fig. 2J,K). Subtracting the average image of AZs expressing the wildtype proteins from those expressing the Unc13A CaM mutant revealed an accumulation of signal closer to the plasma membrane and a reduction of signal farther away (Fig. 2L). Moreover, the vertical line profiles at the AZ center confirmed that the peak of the Unc13A fluorescence was found closer to the plasma membrane and farther away from the BRP C-terminus in the CaM mutant than in the controls (Fig. 2M). Together with our model predictions and physiological measurements, our data are consistent with the CaM domain mutation causing a conformational shift of the MUN domain towards the plasma membrane which increases release site occupancy, enhances baseline transmission, and abolishes STF.

### Mutation of Unc13A’s CaM abolishes acute (minutes’) presynaptic homeostatic potentiation

Regulation of release sites has also been implicated in longer-lasting presynaptic homeostatic plasticity (PHP) (Muller et al., 2012; Weyhersmuller *et al*., 2011). This plasticity mechanism counterbalances the loss of postsynaptic neurotransmitter sensitivity by increasing the amount of AP-induced neurotransmitter release within minutes (Davis and Muller, 2015). Experimentally, PHP can be acutely induced by applying the glutamate receptor antagonist Philanthotoxin (PhTx) (Frank *et al*., 2006). Recent analysis revealed that immediate presynaptic plasticity on the minutes’ timescale can be achieved independently of a structural AZ remodeling needed for long-lasting potentiation, indicating that acute PHP can be achieved with the available presynaptic material (Böhme *et al*., 2019). Thus, we wondered to what extent PHP-similar to STF (Fig. 1)-depended on a rapid enhancement of transmission by increasing the occupation of available release sites and therefore investigated whether mutation of the Unc13A CaM domain might influence PHP (as it occluded STF).

Current clamp recordings from 3^rd^ instar Drosophila laval NMJs (muscle 6, segments A2 & A3) were performed in animals either expressing wildtype Unc13A or the Unc13A CaM^WRWR^ mutant. Prior to recordings, larvae were incubated for 10 min either in standard hemolymph-like (HL3) control solution, or in HL3 solution containing 20 µM PhTx. First, miniature excitatory postsynaptic potentials (mEPSPs) caused by spontaneous synaptic activity were measured in current clamp experiments for 60 s (Fig. 3A, C). Following this, the efferent motoneural axons innervating the same NMJ were stimulated, and the resulting AP-evoked excitatory postsynaptic potentials (eEPSP) were measured (Fig. 3B, D, left panel). Experiments were performed in the presence of low extracellular Ca^2+^ (0.15 mM) to ensure that eEPSPs in Unc13A CaM^WRWR^ mutants were small enough to assume linear summation from mEPSPs (Katz and Miledi, 1979), allowing assessment of the number of SVs released per AP (quantal content, eEPSP amplitude divided by the average mEPSP amplitude of that cell). As expected from its inhibitory action on ionotropic glutamate receptors at the NMJ, application of PhTx potently diminished mEPSP amplitudes (Fig. 3A). Despite this reduced postsynaptic NT sensitivity, AP-evoked eEPSPs were not diminished in control synapses exposed to PhTx, because of a compensatory increase in the quantal content (Fig. 3B), indicating intact PHP. In contrast, NMJs expressing Unc13A bearing the CaM^WRWR^ mutation failed to undergo PHP as PhTx treatment similarly diminished mEPSP and eEPSP amplitudes and no potentiation of the quantal content was detected (Fig. 3 C,D). Thus, enhanced transmission observed in the Unc13A CaM domain mutant similarly occludes PHP on a minutes’ timescale as it abolished STF on a millisecond timescale, indicating that increasing release site occupancy via Unc13A regulatory domains may underlie both forms of presynaptic plasticity.

**Fig. 3:**
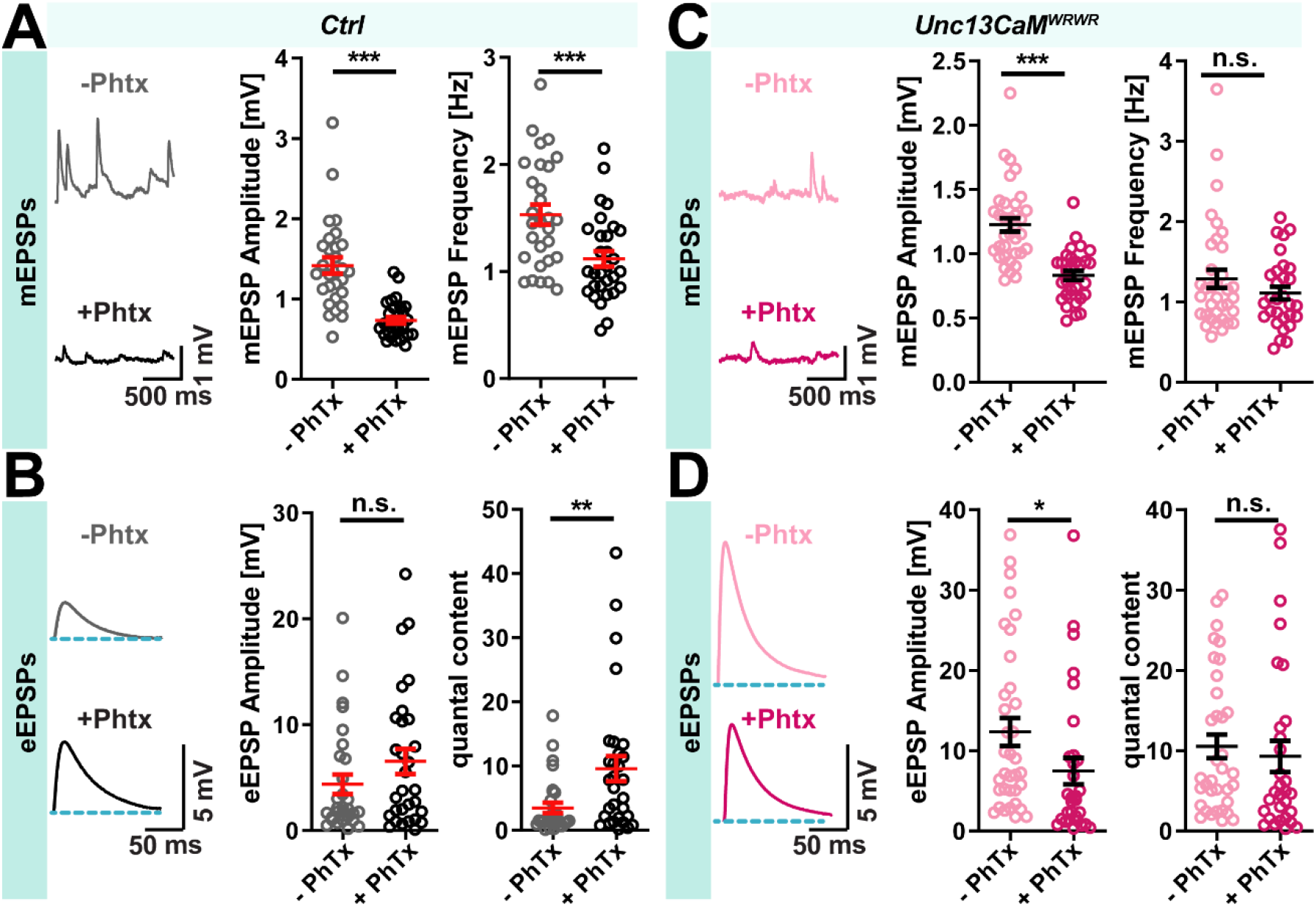
Unc13A CaM^WRWR^ mutation abolishes PhTx-induced PHP. (**A-D**) Graphs showing experimental data acquired in current clamp recordings from larval M6 NMJs at 0.15 mM extracellular Ca^2+^. (**A - D**) Representative example traces from individual cells showing spontaneous activity (**A, C, left panel**) and average traces of AP evoked responses from 5 repetitions (**B, D, left panel**) in in *Ctrl* and *Unc13CaM*^*WRWR*^ NMJs preincubated for 10 minutes with solution containing 20 µM PhTx (+ PhTx) or control solution (containing ddH_2_O of the same volume, PhTx). (**A - D, middle and right panel**) Quantification of spontaneous (**A, C**) and AP evoked transmission (**B, D**) in these experimental conditions (*Ctrl* −PhTx (grey): 29 NMJs from 14 animals, *Ctrl* +PhTx (black): 30 NMJs from 18 animals, *Unc13CaM*^*WRWR*^ −PhTx (light pink): 35 NMJs from 17 animals, *Unc13CaM*^*WRWR*^ +PhTx (magenta): 30 NMJs from 16 animals. (**A, C, middle panel**) PhTx treatment induced similar reduction of mEPSP amplitude in *Ctrl* (**A, middle panel**) and *Unc13CaM*^*WRWR*^ (**C, middle panel**) NMJs. (**A, C, right panel**) Quantification of mEPSP frequency revealed a significant reduction in *Ctrl* synapses (**A, right panel**) and no change in this parameter upon PhTx treatment in *Unc13CaM*^*WRWR*^ mutants (**C, right panel**). (**B, D, middle panel**) Quantification of AP evoked release revealed unaltered amplitudes in *Ctrl* cells (**B, middle panel**), while eEPSP amlitudes were reduced in *Unc13CaM*^*WRWR*^ NJMs upon PhTx treatment (D, **middle panel**). (**B, D, right panel**) Quantal content quantification revealed a significant increase in *Ctrl* NMJs upon PhTx treatment (**B, right panel**), while no change in this parameter could be observed in PhTx treated *Unc13CaM*^*WRWR*^ mutant synapses (**D, right panel**). Genotypes (see methods for details): *Ctrl*: Unc13A and -B null animals expressing wildtype Unc13A; *CaM*^*WRWR*^ mutant: Unc13A and -B null animals expressing the Unc13A *CaM*^WRWR^ mutant. Statistical analysis with Student’s two-tailed t-test. Data shows mean values ± s.e.m., * p ≤ 0.05; ** p ≤ 0.01; *** p ≤ 0.001.

### Structural AZ remodeling-previously linked to long term PHP-persists in Unc13A CaM^WRWR^ mutants

On longer timescales (days) a similar compensatory increase in the quantal content is induced by the genetic deletion of the high conductance glutamate receptor subunit IIA (*GluRIIA*) (Böhme *et al*., 2019; Frank *et al*., 2006; Petersen *et al*., 1997). This long-term PHP-unlike the acute PHP described above-is blocked under conditions preventing structural AZ remodeling (e.g. by mutation of the transport adapter Aplip-1 or the transport-regulating kinase SRPK (Böhme *et al*., 2019; Johnson et al., 2009; Siebert et al., 2015). This is consistent with local enrichments of synaptic proteins (BRP/Unc13A) sustaining PHP. Several treatments have been reported that impede long-term PHP but leave acute PHP intact (Böhme *et al*., 2019; Goel *et al*., 2019; James et al., 2019). However, whether inversely acute PHP is necessary to achieve long-term PHP remains elusive. To investigate whether acute PHP is mandatory for AZ remodeling, we quantified whether PhTx-treatment led to detectable changes in the fluorescence intensities of Unc13A CaM^WRWR^ mutant NMJs stained against BRP and Unc13A. This analysis revealed that despite the occlusion of acute PHP in these mutants, structural AZ remodeling upon PhTx treatment was intact and resulted in increased fluorescence levels in both channels detected by confocal microscopy (Fig. 4B,C). Thus, while release site-based plasticity via the Unc13A regulatory domains is essential for rapid presynaptic plasticity (STF & acute PHP), structural AZ remodeling persists, indicating that long-term PHP might function independently and may still be intact (Fig. 4A).

**Fig. 4:**
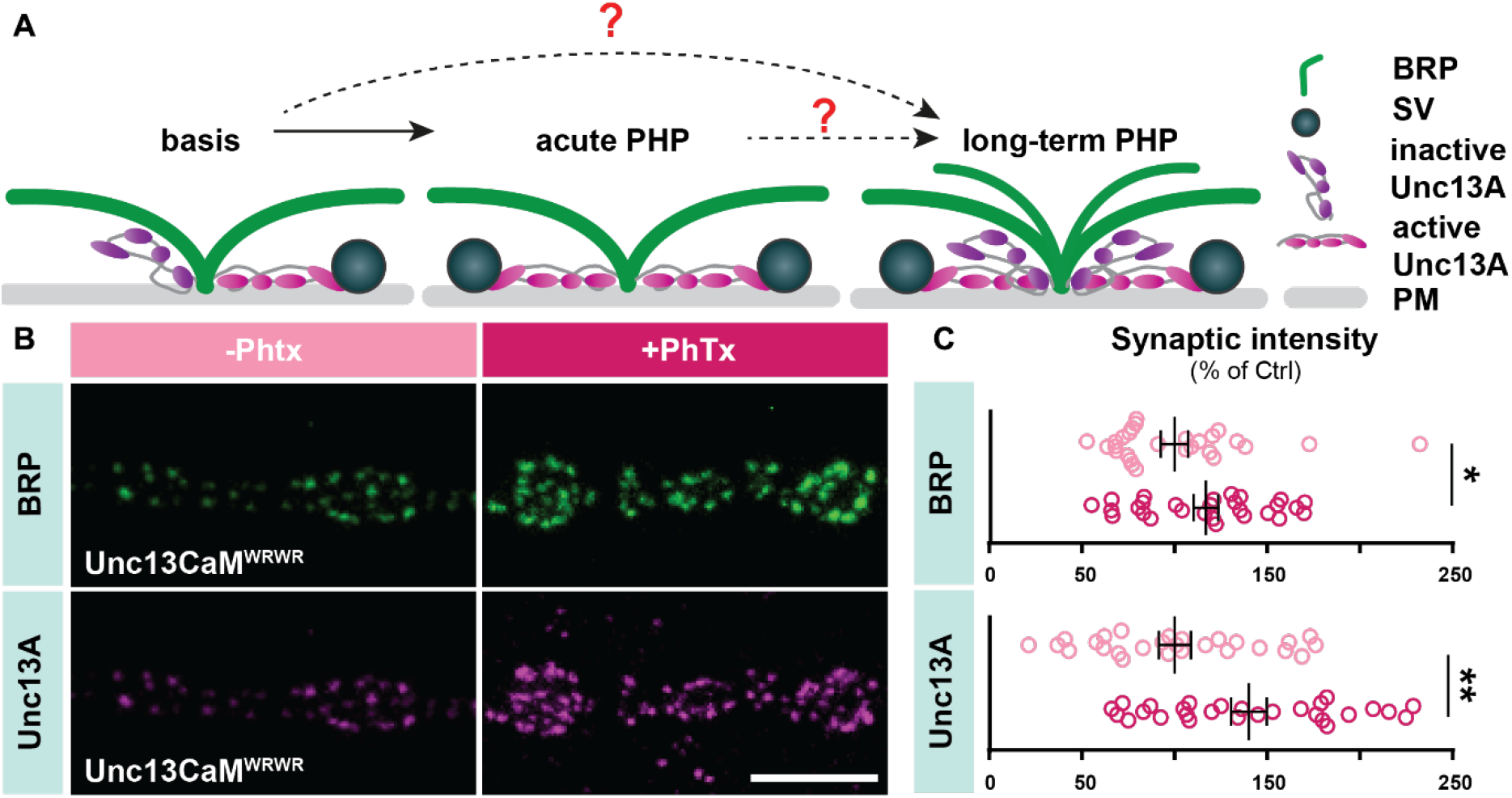
Unc13A CaM mutation does not interfere with PhTx-induced synaptic remodeling. (**A**) Hypothetical scheme of acute and long-term PHP. Acute presynaptic potentiation enhances active release site numbers by rapidly activating available Unc13A while during long-term plasticity scaffolding protein levels (BRP) at the AZ are actively increased which in turn recruits more Unc13A to elevate the total release site number. Whether long-term plasticity can bypass the acute plasticity phase is unknown (red questions marks). (**B**,**C**) Confocal images and quantification of AZ intensities normalized to control condition (−PhTx), of muscle 4 NMJs of abdominal segments 2-4 from 3^rd^ instar larve of *Unc13CaM*^*WRWR*^ mutants labelled with the indicated antibodies without (−PhTx, light pink) and with 10 min 20 µM PhTx (+PhTx, magenta) treatment. Genotype (see methods for details): *CaM*^*WRWR*^ mutant: Unc13A and -B null animals expressing the Unc13A *CaM*^WRWR^ mutant. *Unc13CaM*^*WRWR*^ −PhTx (light pink): 27 NMJs from 6 animals, *Unc13CaM*^*WRWR*^ +PhTx (magenta): 28 NMJs from 6 animals. Scale bar: 5 µm. Statistics: Mann-Whitney U test (BRP) and Student’s t test (Unc13A). * P ≤ 0.05; ** ≤ 0.01. All panels show mean ± s.e.m..

### DAG analogs targeting the Unc13A C1 domain similarly affect baseline transmission and STP

An additional, highly conserved regulatory domain in close proximity of the Unc13A Ca^2+^/Calmodulin binding domain is the Unc13A C1 domain which binds the signaling lipid DAG and the phorbol esters PMA (unnatural DAG analogs) that potently enhance NT release (Basu *et al*., 2007; Betz *et al*., 1998; Maruyama and Brenner, 1991; Rhee *et al*., 2002; Schotten *et al*., 2015; Song et al., 2002). We wondered whether this enhancement depended on a similar activation of release sites (Fig. 1). We first confirmed that acute application of the phorbol ester PMA potentiated synaptic transmission at 3^rd^ instar laval *Drosophila* NMJs expressing wildtype Unc13A (Song *et al*., 2002). For this we used TEVC recordings to measure AP-evoked responses and STP in 0.4 mM external Ca^2+^ under which condition the potentiation of NT release by the Unc13A CaM mutation had been strongest (Fig. 1H,I). This revealed a potent enhancement of synaptic transmission following 10 minutes incubation with 10 µM PMA (Fig. 5A). Quantification of the PPR revealed that PMA treatment furthermore decreased the STF typically observed in control synapses (Fig. 5A). The same analysis performed at more physiological 1.5 mM extracellular Ca^2+^ levels revealed similar consequences but overall smaller effects of PMA-treatment (Fig. S4). These effects are similar to the ones observed upon mutation of the Unc13A CaM binding domain (Figs. 1&5) and we wondered to what extent these two ways of enhancing NT release were interdependent by investigating PMA sensitivity in Unc13A CaM mutants recorded in parallel. This revealed that neither the first eEPSC_1_ amplitudes increased nor the PPRs decreased by PMA treatment in Unc13A CaM mutant animals (Fig. 5E). Thus, mutation of the Unc13A CaM domain occludes further potentiation via the C1 domain, consistent with both treatments inducing maximal release site occupation.

**Fig. 5:**
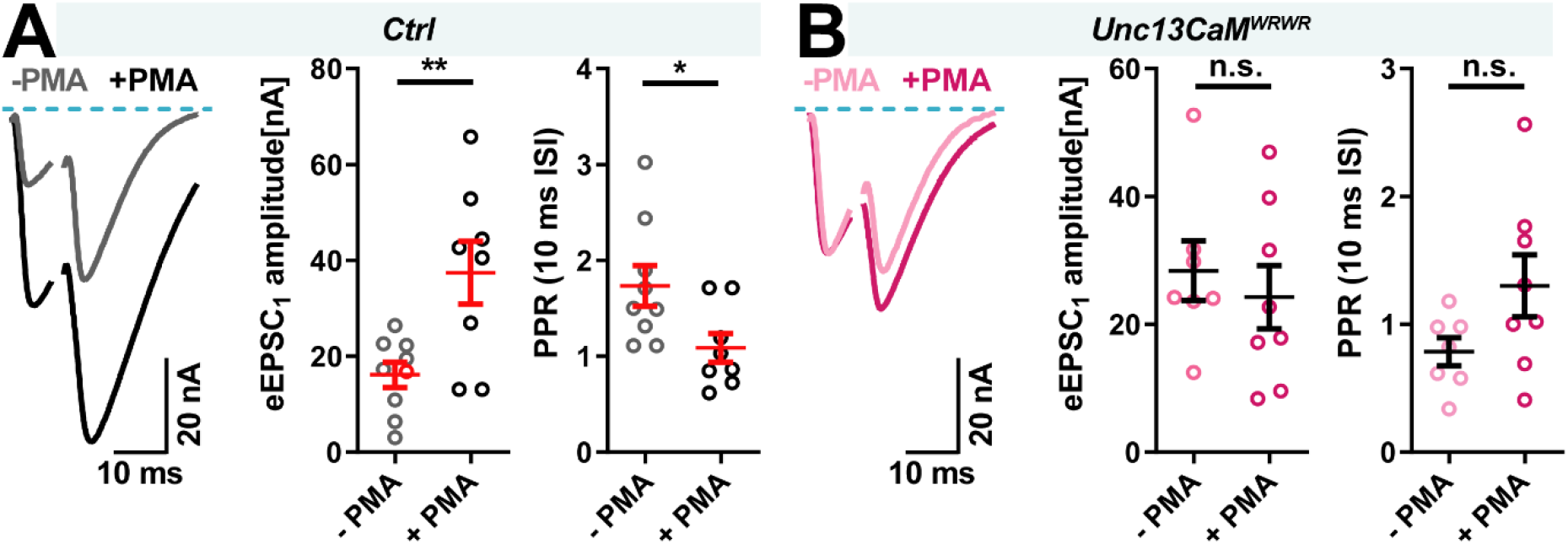
Respective occlusion of presynaptic potentiation by activation of the Unc13A CaM and C1 domains. TEVC experiments from larval M6 NMJs at 0.4 mM extracellular Ca^2+^. (**A**) Analysis of control (*Ctrl*) animals. Left panel: Example traces of AP evoked paired-pulse responses preincubated with DMSO (−PMA) or 2 µM PMA in DMSO. **Middle panel**: Quantification of eEPSC_1_ amplitudes. Right panel: Quantification of PPR ratios (10 ms inter-stimulus interval (ISI)). (**B**) Same analysis as in (A) for *Unc13CaM*^*WRWR*^ mutant flies. Genotypes (see methods for details): *Ctrl*: Unc13A and -B null animals expressing wildtype Unc13A; *CaM*^*WRWR*^ mutant: Unc13A and -B null animals expressing the Unc13A *CaM*^WRWR^ mutant. Number of cells (n) and animals (N) investigated: n(*Ctrl, −PMA*)= 9, N(*Ctrl, −PMA*)= 5; n(*Ctrl, +PMA*)= 8, N(*Ctrl, −PMA*)= 5; n(*Unc13CaM*^*WRWR*^ mutant, *−PMA*)= 9, N(*Unc13CaM*^*WRWR*^ mutant, *−PMA*)= 5; n(*Unc13CaM*^*WRWR*^ mutant, *+PMA*)= 8, N(*Unc13CaM*^*WRWR*^ mutant, *+PMA*)= 5. Dots represent individual cells/NMJs, mean and SEM are shown. Statistical analysis with Student’s two-tailed t-test, p > 0.05: non-significant (n.s.), p ≤ 0.05: *, p≤ 0.01: **.

### Acute application of DAG analogs occludes acute (minutes’) PHP

Due to the functional similarity and non-additivity in the potentiation of NT release caused by targeted manipulation of the Unc13A C1 (PMA application) or CaM^WRWR^ mutation we next wondered whether acute PMA application similarly occluded PhTx-induced PHP as the mutation of CaM domain had (Fig. 4). This was investigated in current clamp NMJ recordings of wildtype (w1118) *Drosophila* 3^rd^ instar larvae (expressing both Unc13A and -B). In control conditions, 3^rd^ instar *Drosophila* larvae were preincubated for 10 minutes without PMA (only the solvent DMSO was added) followed by a second 10-minute incubation in the presence or absence of 20 µM PhTx. As before, PhTx reduced mEPSP amplitudes but did not diminish eEPSP amplitudes due to a compensatory increase in the quantal content, indicating intact PHP (Fig. 6A&B). Wildtype flies incubated with PMA but without PhTx showed similar (in tendency slightly smaller) mEPSP amplitudes as controls without PMA (Fig. 6A,C), but eEPSP amplitudes were markedly increased, demonstrating a PMA-induced increase in the quantal content (Fig. 6B,D). As was the case without PMA, incubation with both PMA and PhTx reduced mEPSP amplitudes compared to PMA-only treated animals (Fig. 6C). Yet unlike without PMA, PhTx reduced average eEPSP amplitudes (statistically not significantly different at a 5% level) and no elevation of the quantal content was seen compared to NMJs treated with PMA alone (Fig. 6D). These findings illustrate that presynaptic potentiation induced by acute PMA treatment occluded PHP to counteract the effect of PhTx, consistent with PMA “maxing out” release site occupation. Together, our findings indicate that acute pharmacological activation of the Unc13A C1 domain (similarly to mutation of its CaM domain) enhances synaptic transmission, and that this enhancement impedes both STF and PHP.

**Fig. 6:**
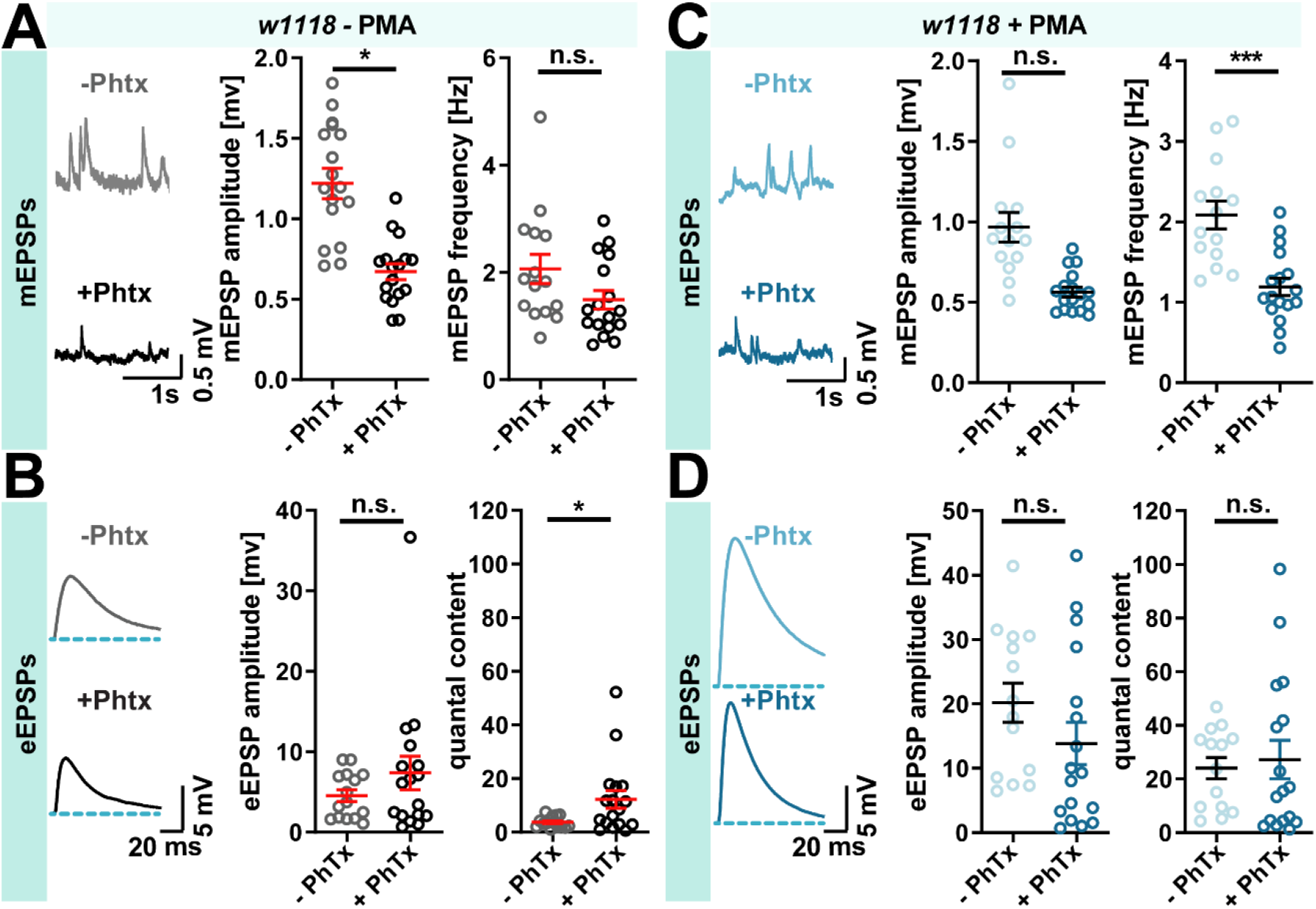
Acute Unc13A-C_1_ activation with PMA blocks PhTx-induced PHP. (**A - D**) Graphs showing experimental data acquired in current clamp recordings from larval M6 NMJs at 0.2 mM extracellular Ca^2+^. (**A - D**) Representative example traces from individual cells showing spontaneous activity (**A, C, left panel**) and average traces of AP evoked responses from 5 repetitions (**B, D, left panel**) in *w*^*1118*^ NMJs pre-incubated for 10 minutes with control solution (−PMA) followed by 10 minutes incubation either with 20 µM PhTx (+PhTx) or without (−PhTx) as well as *w*^*1118*^ NMJs pre-incubated for 10 minutes with 2 µM PMA (+PMA) followed by 10 minutes incubation either with 20 µM PhTx (+PhTx) or without (−PhTx). (**A - D, middle and right panel**) Quantification of spontaneous (**A, C**) and AP evoked transmission (**B, D**) in these experimental conditions (−PMA – −PhTx (grey): 15 NMJs from 8 animals, −PMA +PhTx (black): 17 NMJs from 9 animals, +PMA −PhTx (light blue): 14 NMJs from 7 animals, +PMA +PhTx (blue): 17 NMJs from 9 animals). PhTx treatment induced a similar reduction of mEPSP amplitude upon co-application-with PMA (**C, middle panel**-) as observed in control treatment without PMA (**A, middle panel**-). The mEPSP frequency was reduced upon co-application of PMA and PhTx-(**C, middle panel**) when compared to the PMA only condition (**A, middle panel**), where no significant change of this parameter could be observed. PhTx treatment had no significant effect on eEPSP amplitudes in both the control experiment (**B, middle panel**-) as well as co-application-with PMA (**D, middle panel**-). PhTx treated synapses exhibited a significant increase in quantal content (**B, right panel**-), while preincubation with PMA blocked quantal content increase upon PhTx-treatment (**D, right panel**-). Genotype (see methods for details): *w1118* wildtype. Statistical analysis with Mann-Whitney U test. Mean values ± s.e.m are indicated in plots, * p ≤ 0.05; ** p ≤ 0.01; ** p ≤ 0.001.

## Discussion

Here we investigated putative molecular mechanisms of changes in release site occupation relevant for presynaptic short-term plasticity and homeostatic potentiation of SV release. Our main conclusions are:

- Mutation of the Unc13A CaM domain abolishes STF and increases baseline transmission at low Ca^2+^, consistent with unnaturally high release site occupation;
- The functionally essential MUN domain localizes closer to the plasma membrane in the Unc13A CaM domain mutant, consistent with site occupation being regulated by a conformational switch of Unc13A;
- Mutation of the Unc13A CaM domain abolishes PHP upon acute postsynaptic receptor block, consistent with PHP operating via acute increases of release site occupation;
- Similar effects (enhanced baseline transmission, block of STF and acute PHP) are seen upon pharmacological activation of the Unc13A C1 domain and no further potentiation seen for coincident manipulation, suggesting the same downstream increase in site occupation;
- STF and PHP may thus underlie similar increases in release site occupation due to Unc13A activation;

### Enhanced potentiation of transmitter release revealed at low extracellular Ca^2+^

Analysis of mammalian Munc13-1 and Munc13-2 isoforms early on identified different STP signatures and mutational analysis established a role for the Munc13-1 CaM domain (Junge *et al*., 2004; Rosenmund et al., 2002). The Drosophila Unc13A CaM domain mutant studied here is based on the Munc13-1 CaM mutant and our data confirm its relevance for STP at the *Drosophila* NMJ. Our modelling approach had predicted enhanced effects at low extracellular Ca^2+^ concentrations (Fig. 1D,E) which led us to specifically investigate this condition and revealed strongly elevated initial responses to AP stimulation as one of the hallmarks of this mutation (Fig. 1H,I). Previous analyses were performed at higher levels of extracellular Ca^2+^ where our simulations and experiments also confirm a similar behavior as in wildtype synapses (Junge *et al*., 2004; Lipstein et al., 2013; Ritzau-Jost et al., 2018)(Fig. 1).

### Conformational changes in Unc13A as a mechanism for rapid release site switching

Unc13 proteins have a conserved function in determining presynaptic strength and with their extended structure interact both with AZ scaffolding proteins and SVs (Betz et al., 2001; Bohme *et al*., 2016; Kawabe et al., 2017; Padmanarayana et al., 2021; Quade et al., 2019; Reddy-Alla *et al*., 2017; Sakamoto *et al*., 2018; Wang et al., 2016; Xu et al., 2017; Zhou *et al*., 2013). Consequently, conformational changes of (M)Unc13 have been proposed to regulate NT release and STP (Camacho *et al*., 2021; Grushin *et al*., 2022). Because mutation of the Unc13A CaM domain resulted in effects consistent with an increase of release site occupation (elevated baseline transmission and loss of STF at low extracellular Ca^2+^, Fig. 1), we investigated possible changes in the orientation of its functionally essential MUN domain (Basu *et al*., 2005; Stevens *et al*., 2005) (Fig. 2). This revealed a slight shift of the MUN domain towards the plasma membrane in Unc13A CaM^WRWR^ mutants while its distance to the AZ center (and the overall fluorescence per AZ) was not affected (Figs. 2&S2). Our data are thus both consistent with propositions that closer membrane localization increases SV priming (Camacho *et al*., 2021; Grushin *et al*., 2022) and that this is dynamically increased during STF (Kobbersmed *et al*., 2020; Lin *et al*., 2022; Silva *et al*., 2021) while the CaM^WRWR^ mutation of Unc13A “locks” the protein in this high activity sate promoting release site occupancy.

On the other hand, this redistribution towards the plasma membrane likely makes it more difficult for to stabilize vesicles at intermediate distances usually captured by the extended Unc13A conformation. If these vesicles serve to repopulate release sites during repetitive activation, this could explain why inhibiting (M)Unc13-calmodulin interaction impairs the replenishment of readily releasable vesicles after strong stimulation at the murine calyx of Held synapse (Lipstein *et al*., 2013; Sakaba and Neher, 2001).

### Different phases of PHP by cooperating mechanisms

Classical experiments demonstrated robust PHP in current clamp recordings performed at low extracellular Ca^2+^ (Böhme *et al*., 2019; Frank *et al*., 2006). In these experiments, low Ca^2+^ was chosen for the technical reason that the quantal content can only be assessed when APs induce small eEPSPs as otherwise these do not sum linearly from the mEPSPs recorded at rest (Katz and Miledi, 1979). Moreover, our short-term plasticity model predicts high capacity for site-based plasticity at low extracellular Ca^2+^ (∼240%, ∼220% and 150% at 0.4, 0.75 and 1.5 mM external Ca^2+^) due to a low steady state release site occupation (41.1%, 45.0% and 66.7%)(Fig. S1) (Kobbersmed *et al*., 2020). However, these estimates should be taken with caution as they rely entirely on parameters previously estimated from NMJ’s STP behavior (Kobbersmed *et al*., 2020) and additional aspects not captured by the model may be relevant for PHP. On the other hand, as increased release site occupation appears to be possible during millisecond STF (Fig. 1B_1_,B_2_), PHP might be much faster than previously thought and could correspond to the “rapid, low-gain” homeostatic response on a timescale of seconds (Frank *et al*., 2006).

Following this immediate “ultrafast PHP”, further compensatory reactions appear necessary to reset the functional state of the synapse. Remarkably, when assayed 10 min after PhTx treatment, potentiated synapses have the same baseline STP profile as non-potentiated ones and TEVC recordings at this time demonstrated PHP even at an extracellular Ca^2+^ concentration of 15 mM (Muller et al., 2015; Ortega et al., 2018). On this extended timescale, changes of the AZ composition are observed that appear required for long-term stabilization of potentiation (Böhme *et al*., 2019; Mrestani et al., 2021; Turrel et al., 2021). And while this remodeling is dispensable for PHP 10 min after PhTx treatment in mutants where this is blocked, it is currently unknown whether the transition to stable PHP can be faster when remodeling is intact or in cases where “ultrafast PHP” is exhausted (e.g. at high extracellular Ca^2+^). On the other hand, synaptic remodeling *alone* is clearly insufficient for potentiation, because we show that 10 min after PhTx treatment synaptic remodeling is intact in Unc13A CaM^WRWR^ mutants while PHP at this time is blocked (Fig. 5). Thus, an important question future studies should address is how long-term potentiation achieves a stable reset of the physiological properties from the “ultra-rapid” PHP we describe here.

### Unc13A regulatory domains as an integration hub for release site-based plasticity

Auto-inhibitory functions were described for regions spanning the (M)Unc13 C2 (C2A & C2B) and C1 domains (Camacho *et al*., 2017; Deng et al., 2011; Li et al., 2019; Michelassi *et al*., 2017), and our findings indicate that this further extends to its CaM domain. Binding of signaling molecules to these domains could relieve this inhibition and support release site occupation. The variety of Ca^2+^- and lipid signals sensed by these domains could well reflect the physiological regulation of diverse forms of synaptic plasticity integrating on the release sites. While our findings indicate the downstream activation of Unc13A is relevant for STF as well as for PHP, our experiments do not allow us to identify which signals are natively responsible for these plasticity phenomena. For instance, while the mutation of the Unc13A CaM domain could induce its effects by impairing interactions with Ca^2+^/Calmodulin (Junge *et al*., 2004), it could also directly render Unc13A hyperactive by inducing the conformational/functional change which bypasses the need for the native activation. Consistent with the latter option, the same qualitative effects (increased baseline transmission and loss of STF at low extracellular Ca^2+^ and a loss of PhTx-induced PHP) were observed upon acute PMA application targeting the Unc13A C1 domain (Figs. 5&6) and the effects were non-additive (Fig. 5B). This suggests that manipulation of either domain similarly and maximally increases release site occupation. Notably, mutation of the Munc13-1 CaM domain did not interfere with the potentiation of synaptic transmission in mouse hippocampal neurons upon treatment with another phorbol ester, PdBu (Junge *et al*., 2004). These differences could be due to the different treatment conditions (duration of treatment, extracellular Ca^2+^ concentration (Junge *et al*., 2004)) or different actions of the two compounds (as observed in other systems (Gaudry et al., 1990)). Another yet unexplored possibility is that differences in the local lipid environment in these systems might cause different sensitivities to these compounds.

Our model predicts a Ca^2+^-dependent change in release site occupation for STF, but whether this is mediated by direct Ca^2+^-binding to one of Unc13As regulatory domains (e.g. Ca^2+^/CaM, Ca^2+^/C2) (Junge *et al*., 2004; Shin *et al*., 2010) or involves additional components of the release machinery is not known. Notably, loss of the Ca^2+^ sensing protein synaptotagmin-7, which also binds Calmodulin and functions in STF, SV replenishment and vesicle (re)docking (Jackman et al., 2016; Liu et al., 2014; Tawfik et al., 2021; Wu et al., 2022), similarly increased baseline transmission and abolished STF at low Ca^2+^ as we observe here (Figs. 1, 5) (Fujii et al., 2021; Guan et al., 2020). This could indicate a cooperative function with Unc13A for STF. In addition, the Ca^2+^ sensing protein Doc2 interacts with Munc13-1 and affects its subcellular localization which could affect release site occupation on distinct timescales and contribute to Doc2’s role in asynchronous/spontaneous NT release or synaptic augmentation (Groffen et al., 2010; Mochida et al., 1998; Orita et al., 1997; Xue et al., 2018; Yao et al., 2011). On longer timescales the ER-resident Ca^2+^ sensing protein MCTP is relevant for PHP (Genc et al., 2017), but whether this directly affects Unc13A or involves additional messengers (e.g. DAG/PI(4,5)P_2_) is unknown. Nevertheless, a picture is emerging in which Unc13A proteins are highly relevant plasticity targets that respond to a multitude of signals (Ca^2+^, Ca^2+^/Calmodulin, DAG and PI(4,5)P_2_/Ca^2+^) to regulate release site participation. The broad range of signals integrated likely relates to different forms of synaptic plasticity (e.g. facilitation/augmentation/potentiation) across extended timescales that all converge on Unc13A proteins for release site based plascitiy.

## Methods

### Mathematical modelling and simulation

We performed model simulations to investigate the effects of hypothetical mutants that make vesicle unpriming insensitive to Ca^2+^. For this purpose, we used a previously published model of synaptic release describing the priming, unpriming and fusion of vesicles as a function of Ca^2+^ (Kobbersmed *et al*., 2020). In brief the model works as follows:

First, for various extracellular Ca^2+^ concentrations (0.4mM, 0.75mM, 1.5mM, 3mM, 6mM) and for different locations from the Ca^2+^ sources (location 120 nm was used to plot Fig. B_1_ and Fig. C_1_), Ca^2+^ transients induced by two APs (10 ms interstimulus interval) were determined. This was done by deterministic simulation using the Calcium Calculator program, version 6.8.6 (CalC 6.8.6) created by Victor Matveev (Matveev et al., 2002). Next, 180 release sites were placed at heterogeneously located positions from the Ca^2+^ source. The Ca^2+^ transient corresponding to each release site was read out as the input for the exocytosis simulation. This simulation stochastically computes vesicle priming, unpriming and fusion of primed vesicles. Synaptic vesicle fusion was modelled following the single-sensor model designed by (Lou et al., 2005), and thus depended on internal Ca^2+^ concentration. Furthermore, the internal Ca^2+^ concentration also affected the unpriming rate (see below). The stochastic simulations were repeated 200 times for each extracellular Ca^2+^ concentration. From the stochastically simulated fusion times, eEPScs were generated using convolution with a mEJC. The amplitude of the second eEPSC was determined after subtraction of the extrapolated decay of the first eEPSC. The paired-pulse ratio (PPR) corresponds to the ratio between the amplitude of the second EPSC (determined as above) and the first (PPR = eEPSC_2_ / eEPSC_1_).

For our study, we wanted to change the Ca^2+^-dependent unpriming rate in the model, which is calculated using the following mathematical equation:

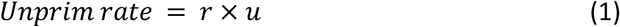

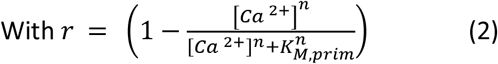

In this equation, [Ca^2+^] is the intracellular Ca^2+^ concentration, *u* is the basal unpriming rate constant, *K*_*M, prim*_ is a Michaelis-Menten-like constant, and *n* is the Ca^2+^ cooperativity. The speed of unpriming (occurring at basal rate u) is slowed by elevations in Ca^2+^ (r decreases as Ca^2+^ increases, see Fig. S1A). The intracellular Ca^2+^ concentration at rest could be computed from the extracellular Ca^2+^ concentration using the following formula:

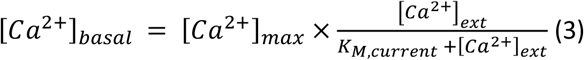

In this equation, [Ca^2+^]_basal_ is the resting intracellular Ca^2+^ concentration, [Ca^2+^]max is the maximal intracellular Ca^2+^ concentration ([Ca^2+^]_max_ = 190nm), [Ca^2+^]_ext_ is the extracellular Ca^2+^ concentration and K_M,current_ is the extracellular Ca^2+^ concentration corresponding to the half of maximal intracellular Ca^2+^ concentration (see Kobbersmed et al., 2020)

To simulate WT condition, we used the same parameters as published for the Ca^2+^ unpriming model by (u = 236.82 s^−1^, n =5, K_M,prim_= 55.21 nM^−1^) (Kobbersmed *et al*., 2020). For the “All sites *off*” r was set to one for all Ca^2+^ concentrations, making the unpriming rate of vesicles constantly high. For the “All sites on” mutant, r was set to zero for all Ca^2+^ concentration. This means that primed vesicles in this mutant condition are unable to unprime. All these simulations were performed on MATLAB (version R2017b).

### Fly Husbandry, Stocks, and Handling

Fly strains were reared under standard laboratory conditions (Sigrist et al., 2003) and raised at room temperature (Bloomington recipe). For experiments both male and female third instar larvae (larval stage 3) were used. The following genotypes were used: Wild-type: +/+ (*w*^*1118*^) (obtained from Bloomington Drosophila Stock Center), *CaM*^*W1620R,W1622R*^:

*;;CaM*^*W1620R,W1622R*^*/CaM*^*W1620R,W1622R*^*;P84200/P84200* (Reddy-Alla *et al*., 2017), *Control:*

*;;Unc13B*^*null*^*/Unc13B*^*null*^*;P84200/ P84200* (Bohme *et al*., 2016; Reddy-Alla *et al*., 2017).

### Electrophysiology: Setup and data acquisition

For all electrophysiological experiments, third instar larvae were dissected on Sylgard (184, Dow Corning, Midland, MI, USA) in Ca^2+^free HL3, (put composition here, pH adjusted to 7.2 (Stewart et al., 1994)) or modified low Mg^2+^ HL3 (pH adjusted to 7.2 (Böhme *et al*., 2019): 70 mM NaCl, 5 mM KCl, 10 mM MgCl_2_, 10 mM NaHCO_3_, 5 mM Trehalose, 115 mM D-Saccharose, 5 mM HEPES) as described previously (Jan and Jan, 1976; Zhang and Stewart, 2010) and transferred to the recording chamber containing HL3 solution with Ca^2+^.

Electrophysiological current clamp and TEVC recordings were performed at 21°C from muscle 6 of abdominal segments A2 and A3 using sharp glass electrodes (borosilicate glass with filament, 0.86 × 1.5 × 80 nm, Science products, Hofheim, Germany) pulled with a P97 pipette puller (Sutter Instrument, CA, USA). Pipettes were backfilled with 3 mM KCl solution and exhibited resistances between 20 to 30 MΩ. Signals were recorded using a 5 KHz lowpass filter at a sampling frequency of 20 kHz using the Digidata 1440A digitizer (Molecular devices, Sunnyvale, CA, USA) with Clampex (v10.6) software and an Axoclamp 900A amplifier (Axon intstruments, Union City, CA, USA) with Axoclamp software. Cells with resting membrane potentials higher than −49 mV and resistances below 4 MΩ prior to measurements were excluded from the datasets. For all TEVC recordings, the postsynaptic muscle cell was clamped at −70 mV and only cells with absolute leak currents below 10 nA throughout individual experiments were included into the analysis. Obtained data was analyzed using Clampfit software (v10.6.2.2). All graphs show mean values ± S.E.M and were generated with GraphPad Prism 6.01 and Adobe Illustrator (Adobe Systems, San Jose, CA, USA).

### Baseline transmission and STP in Unc13ACaM mutants

For experiments shown in **Fig. 1** TEVC recordings were performed using normal HL3 solution (pH adjusted to 7.2 (Stewart *et al*., 1994)). Cells were stimulated following a paired-pulse paradigm by giving two short (300 µs) depolarizing pulses of 9 V in short succession (10 ms interstimulus interval) 10 times (0.1 Hz) each at increasing (0.4, 0.75, 1.5, 3 and 6 mM) extracellular Ca^2+^ concentrations and eEPSCs were recorded. Cells were stimulated following a paired-pulse paradigm by giving two short (300 µs) depolarizing pulses of 9 V in short succession (10 ms interstimulus interval) 10 times (0.1 Hz) each at increasing (0.4, 0.75, 1.5, 3 and 6 mM) extracellular Ca^2+^ concentrations and eEPSCs were recorded. Successive increase of the extracellular Ca^2+^ concentration was achieved by exchanging 1 ml of the bath solution (total volume: 2 ml) with HL3 of a higher calcium concentration (exchange solution, see Table 1) and slowly mixing them in the bath using the pipette. After each titration step the larvae were acclimated in the recording chamber for 1 minute prior to recording. EPSCs and PPRs shown in **Fig. 1** were analyzed using a custom-written MATLAB script to correct for residual current caused by the preceding eEPSC_1_ as reported previously (Kobbersmed *et al*., 2020). PPRs were calculated by dividing the amplitude of eEPSC_2_ by eEPSC_1_ and averaging these values for 10 stimulations in each cell.

**Table 1:** Ca^2+^ concentrations of titration solutions.

| Ca<sup>2+</sup>-concentration of exchange solution<br>(1 ml exchange solution added to 1 ml solution in<br>recording chamber) | Total Ca<sup>2+</sup>-concentration in recording<br>chamber<br>(total volume: 2 ml) |
| --- | --- |
| - | 0.4 mM |
| 1.1 mM | 0.75 mM |
| 2.25 mM | 1.5 mM |
| 4.5 mM | 3 mM |
| 9 mM | 6 mM |

### Rapid PHP in Unc13-CaM mutants

Electrophysiology: For experiments shown in **Fig. 3&6**, current clamp experiments were performed using low Mg^2+^ HL3 (pH adjusted to 7.2 (Böhme *et al*., 2019): 70 mM NaCl, 5 mM KCl, 10 mM MgCl_2_, 10 mM NaHCO_3_, 5 mM Trehalose, 115 mM DSaccharose, 5 mM HEPES). Rapid PHP was induced using a final concentration of 20 µM of the glutamate receptor blocker Philantotoxin433 (PhTx433, SigmaAldrich, MO, USA, stored at 20°C as 4 mM stock in double distilled water (ddH2O)). The same volume of ddH_2_O was used in control experiments. Incubation in 30 µl Ca^2+^free low Mg^2+^ HL3 +/ PhTx (10 minutes) and dissection were performed as reported previously (Böhme *et al*., 2019). The specimen were rinsed 3 times with PhTx and Ca^2+^free low Mg^2+^ HL3 and transferred into the recording chamber. Because the evoked amplitudes are particularly high in Unc13A *CaM*^*WR,WR*^ mutants such that the assumption of linear summation of mEPSPs to eEPSPs is no longer valid, these recordings were performed at 0.15 mM extracellular Ca^2+^ (Bykhovskaia and Vasin, 2017).

### Assessment of phorbol ester induced presynaptic potentiation in Unc13-CaM mutants

For experiments shown in **Figs. 5** and **S4**, TEVC experiments were performed using normal HL3 (pH adjusted to 7.2 (Stewart *et al*., 1994)). Prior to dissection, animals were pre-incubated for 10 minutes in HL3 (0 mM Ca^2+^) containing 2 µM of phorbol ester Phorbol-12-myristat-13-acetat (PMA (Sigma Aldrich, Germany), stored at −20°C as 10 mM stock in dimethyl sulfoxide (DMSO)), or the same volume of DMSO for control experiments. For this, a vertical incision to the larval body wall muscles was made and 30 µl of the incubation solution was pipetted into the abdominal larval cavity. During the last 2 minutes of incubation, larvae were dissected, rinsed 3 times with Ca^2+^-free HL3 once incubation was completed and transferred into the recording chamber containing HL3 (containing 0.4 mM or 1.5 mM Ca^2+^). The postsynaptic muscle cell was clamped at −70 mV in TEVC mode and only cells with absolute leak currents below 10 nA throughout recordings were included into the analysis. Cells were stimulated 10 times by giving two paired stimulating pulses to the innervating motoneuron (300 µs pulses of 8 V, paired pulse stimulation with 10-ms-inter-stimulus interval repeated at 0.1 Hz). Analysis of evoked was performed using pClamp software (Molecular devices, Sunnyvale, CA, USA) by estimating the amplitudes for eEPSC_1_ and eEPSC_2_ for each stimulation and averaging these to generate cell averages. Paired pulse ratios (PPRs) were calculated by dividing the amplitude of eEPSC_2_ by eEPSC_1_ and averaging these values for 10 stimulations in each cell.

### Phorbol ester induced presynaptic potentiation in rapid PHP

For data presented in **Fig. 6**, current clamp experiments were performed using low Mg^2+^ HL3 (pH adjusted to 7.2 (Böhme *et al*., 2019): 70 mM NaCl, 5 mM KCl, 10 mM MgCl_2_, 10 mM NaHCO_3_, 5 mM Trehalose, 115 mM DSaccharose-, 5 mM HEPES). Prior to dissection, animals were first pre-incubated for 10 minutes in low Mg^2+^ HL3 (10 mM Mg^2+^, 0 mM Ca^2+^) containing 2 µM of phorbol ester Phorbol-12-myristat-13-acetat (PMA (Sigma Aldrich, Germany), stored at −20°C as 10 mM stock in DMSO) or the same volume of DMSO for control experiments. For this, a vertical incision to the larval body wall muscles was made and 30 µl of the incubation solution was pipetted into the abdominal larval cavity. After 10 minutes, the first incubation solution was removed exchanged by 30 µl of low Mg^2+^ HL3 (0 mM Ca^2+^, 10 mM Mg^2+^) containing PhTx (20 µM) or the same volume of ddH_2_O in control treatments and incubated for further 10 minutes. After preparation, the larvae were rinsed 3 times with Ca^2+^-free low Mg^2+^ HL3 (10 mM Mg^2+^) transferred into the recording chamber containing 0.2 mM Ca^2+^. This concentration of extracellular Ca^2+^ was chosen based on a baseline experiment where eEPSP responses after PMA treatment were probed (not shown in this paper), so that a ceiling effect of evoked responses could be avoided and the QC could be determined accurately (Bykhovskaia and Vasin, 2017). mEPSPs were recorded for 60 s prior to evoking eEPSP. eEPSP responses were elicited by giving 5 short stimulation pulses (300 µs, 8 V, 0.1 Hz) to the respective nerve. For analysis, mEPSP traces were post hoc filtered with a 500 Hz Gaussian lowpass software filter and mEPSP templates were generated for each cell. Using this template, events were identified throughout the 60 s trace and used for analysis. eEPSP amplitudes were calculated by averaging all 5 stimulated responses and quantal contents were calculated by dividing average eEPSP by the average mEPSP-amplitude in each cell.

### Immunostaining, confocal/STED microscopy, image processing and analysis

Third-instar larvae were placed onto a dissection plate with its dorsal side facing up. Larvae were fixed by placing fine pins in the head and the tail (Austerlitz INSECT PINS; 0.10mm). Afterwards 50 μl ice-cold modified hemolymph-like solution (HL3, pH adjusted to 7.2 ((Stewart *et al*., 1994): 70 mM NaCl, 5 mM KCl, 20 mM MgCl2, 10 mM NaHCO3, 5 mM Trehalose, 115 mM D-Saccharose, 5 mM HEPES) was pipetted onto the larvae. Sharp dissection scissors were used to make a small incision in the dorsal, posterior midline of each larva. From that cut on larvae were then completely opened along the dorsal midline to its anterior end. If required, rapid pharmacological homeostatic challenge was assessed by incubating these semi-intact preparations in 20 µM PhTx diluted in Ca^2+^-free, low Mg^2+^ HL3 (see above) for 10 min at room temperature. Controls were treated in the same way but were incubated in HL3 containing a similar amount of dH2O instead of PhTx for 10 min. Following this, the epidermis was pinned down twice on each side and slightly stretched. For rapid pharmacological homeostatic challenge, extreme care was taken to avoid excessive stretching of body wall muscles, as this may significantly impair induction of homeostasis. The internal organs and tissues were removed carefully with forceps. Subsequently, the dissected larvae were washed 3x with 50 μl ice-cold HL3 on the dissection plate and afterwards fixed for 5 minutes with ice-cold methanol. For rapid pharmacological homeostatic challenge the preparation was directly rinsed three times with ice-cold methanol without a previous HL3 washing step. After fixation, larvae were rinsed 3x with ice-cold HL3 on the pad and then transferred into a 5% native goat serum (NGS; Sigma-Aldrich, MO, USA, S2007) diluted in phosphate buffered saline (Carl Roth, Germany) with 0.05% Triton-X100 (PBT). The sample was blocked for 1h at room temperature and then incubated with primary antibodies mouse Nc82 = anti-BRP^C-term^ (1:200 for confocal and 1:500 for STED microscopy, Developmental Studies Hybridoma Bank, University of Iowa, Iowa City, IA; AB Registry ID: AB_2314865) and guinea-pig Unc13A (Figure 4, 1:500;(Bohme *et al*., 2016)) or rabbit Unc13-C-Term (Figure 2, 1:100;(Reddy-Alla *et al*., 2017)) diluted in 5% NGS in PBT overnight at 4 degrees. The next day, the sample was briefly rinsed 5x with PBT and then washed again 4x in PBTP for 20 minutes. Afterwards, the larvae were incubated for 4h with fluorescence labeled secondary antibodies. For confocal imaging: goat anti guinea pig Alexa-Fluor-488 (1:500, Life Technologies A11073, CA, USA), goat anti-mouse-Cy3 (1:500, Jackson ImmunoResearch 115-165-146) and goat anti-HRP-647 (1:500, Jackson ImmunoResearch 123-605-021, PA, USA). For STED microscopy: anti-rabbit Star Red (1:100, Abberior STRED-1001-500UG) and anti-mouse Star Orange (1:100, Abberior STORANGE-1002-500UG). All secondary antibodies were diluted in 5% NGS in PBT. Afterwards, the samples were briefly washed 3x with PBT and then washed again 4x in PBT for 15 minutes. For confocal imaging larvae were mounted in Vectashield (Vector labs, CA, USA) and for STED imaging in ProLong™ Gold Antifade Mountant (Thermo Fisher P36930) on a microscope slide (Carl Roth, Germany; H868) and sealed with a coverslip (Carl Roth, Germany, H 875).

Confocal microscopy was performed with a Leica SP8 microscope (Leica Microsystems, Germany). Images of fixed and live samples were acquired at room temperature. Confocal imaging of NMJs was done using a z-step of 0.25 μm. The following objective was used: 63×1.4 NA oil immersion for NMJ confocal imaging. All confocal images were acquired using the LAS X software (Leica Microsystems, Germany). Images from fixed samples were taken from muscle 4 of 3rd instar larval 1b NMJs (segments A2-A5). Confocal stacks were processed with ImageJ software (http://rsbweb.nih.gov/ij/). Quantifications of AZ protein levels (scored via BRP) were performed as previously described (Andlauer and Sigrist, 2012; Böhme *et al*., 2019). Briefly, the signal of the HRP-647 antibody was used as template for a mask, restricting the quantified area to the shape of the NMJ. To determine the synaptic protein levels, a custom-written ImageJ script was used that detects the locations with highest local maxima in pixel values of the BRP image to generate regions of interests (ROIs) and sets a point selection at each. The intensities were then measured and all selections were deleted, leaving intensity values and (x,y) locations in the results list. The results list was then used to create a circle of size = 5 pixels (pixel size 100 nm) centered around each (x,y) location and the integrated density within these ROIs was measured and taken for further calculations. The same ROIs were then used in the Unc13A channel to measure synaptic Unc13A levels accordingly. Images for figures were processed with ImageJ software to enhance brightness using the brightness/contrast function.

An Abberior Infinity Line confocal and 3D STED super-resolution microscope was used to record two-color STED images. The microscope combines four pairs of excitation laser beams of 640 nm, 561nm, 485nm and 405 nm wavelength with one STED fiber laser beam at 775nm. For this analysis, the excitation lasers 640 nm and 561 nm were used together with the STED laser. 2D STED was performed on a single confocal section. The pixel size was set to 20 nm and the power of the excitation and STED laser chosen such that we gained the highest resolution with least background noise and kept for all NMJs imaged on one day. Imspector Software 16.3.14287-w2129 (Max Planck Innovation, Germany) was used to acquire all STED images. For figures the images were processed with ImageJ software 1.53c to enhance brightness/contrast.

Heatmaps and intensity/line profiles (Fig.2) were generated from co-stained STED images. BRP was stained together with Unc13A C-term. The BRP signal was used to align the Unc13A signal, similar to a previous analysis (Reddy-Alla *et al*., 2017). First, laterally viewed AZ were cut out in both channels (size 25×25, pixel size 20 nm). Then, we manually drew line ROIs through the center of the elongated BRP structure in all individual images (first channel). The lines were placed such that they followed the longitudinal section of the BRP signal/structure. The lines were saved in a third channel. To indicate the location of the plasma membrane lines were added facing towards the cytoplasm and saved as a fourth channel. A custom-written MATLAB (2022a) script was used to align all AZs to the middle and rotated such that all AZs face towards the plasma membrane. In this script, we first determined the midpoint of the line ROIs (third channel) by taking the average of the coordinates of the start and end points of the line. The computed midpoint was rounded to whole integer values. The images in the BRP and Unc13 channel, as well as the line ROI itself and the direction line in the fourth channel, were translated by the difference between the computed midpoint of the line and the desired midpoint of the line (coordinate (13,13)). Subsequently, we rotated the images in the four channels such that the line ROIs would be orientated horizontally. For this we first computed the current angle of the line ROI with the horizontal midline using:

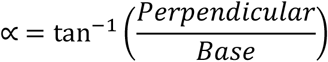

The perpendicular and base line were computed by taking the difference between y-coordinates and x-coordinates, respectively, of the start and end points of the translated line ROI. All four channels were rotated with this angle using the MATLAB function imrotate. As a final step in the rotation of the images, we rotated all channels by 180 degrees when the direction line ROI in the fourth channel was below the horizontal midline in the image. This generates images in which the plasma membrane is facing downwards.

Individual images were normalized such that the total value of all elements in the 25×25 image summed to 1. Afterwards, average images were computed.

To generate the line profiles showed in Figure 2K-L, we extracted the intensity values in the BRP and Unc13 channels of the vertical line through the midpoint of the image. For each individual AZ, we scaled these intensity values such that the area under the curve would sum to 1. The line profiles in figure 2K-L show the average of these scaled intensity lines.

To generate the average images and line profiles in Fig S3, we followed a similar procedure. In this case, we, however, drew circles on the top view BRP images and aligned images from both the BRP and the Unc13 channel to the midpoint of this circle.

## Acknowledgements

This work was supported by grants from the Deutsche Forschungsgemeinschaft to A.M.W. (Emmy Noether Programme, Project Number 261020751 and the TRR 186, Project Number 278001972) and the Novo Nordisk Foundation (Young Investigator Award NNF19OC0056047 to A.M.W.).

