## Supplementary figures for "Release site plasticity via Unc13A regulatory domains mediates synaptic short-term facilitation and homeostatic potentiation"

Supplementary material

### Supplementary figures

Fig. S1

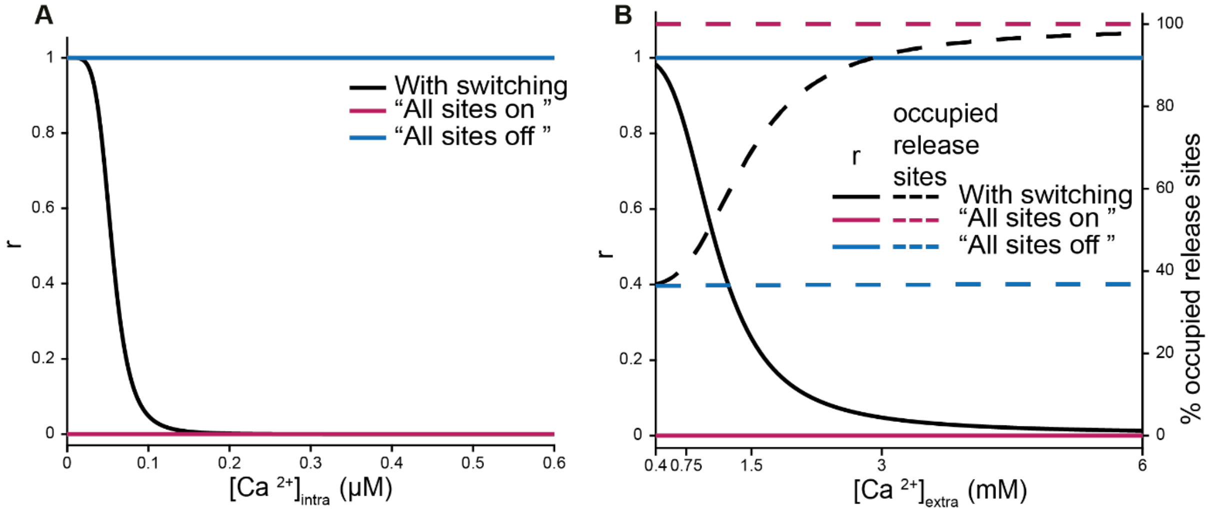


**Fig. S1 Calcium dependence of SV unpriming for simulations with dynamic release site switching and when all sites are constantly switched "on" or "off".** (**A**) The value of r, which scales basal vesicle unpriming rate (u, u*r), as a function of internal [Ca $^{2+}$] (see methods). The "All sites *on*" and "All sites *off*" conditions have respectively a minimum and maximum value for r reflecting a low and high unpriming rate. The black curve refers to the control condition which has release site switching, the pink one to the "All sites *on*" and the blue one, "All sites *off*". (**B**) The value of r as a function of external [Ca $^{2+}$] (solid lines, left axis, see methods) and the percentage of occupied release sites at rest (dashed lines, right axis), depending on external [Ca $^{2+}$].

**Fig. S2**

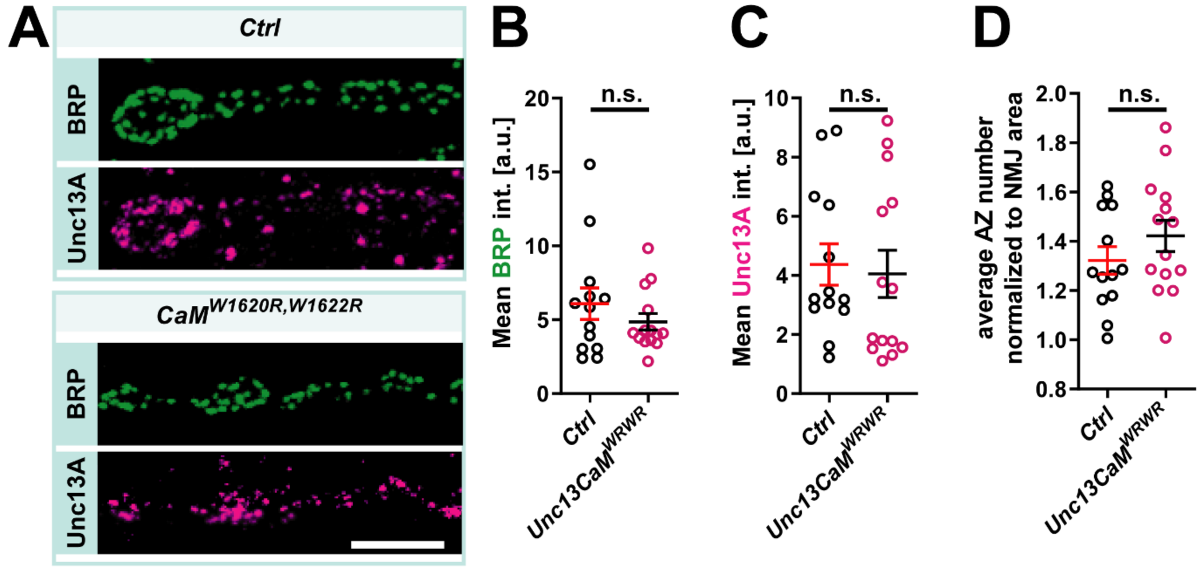


**Fig. S2: CaM binding site mutation does not affect presynaptic levels of Bruchpilot or Unc13A.** Analysis using immunohistochemistry and confocal microscopy at M4 NMJs. (**A**) Example images for Bruchpilot (BRP, green) and Unc13A (magenta) staining in *Ctrl* and *Unc13CaM*$^{WRWR}$ mutant flies. (**B - D**) Quantification of mean per-AZ intensity for BRP (B), Unc13A (C) and mean AZ density (D) averaged per NMJ revealed no major difference between genotypes. Values depicted are mean AZ intensity per NMJ, shown for all NMJs investigated (individual data points) together with mean their mean and SEM. Genotypes (see methods for details): *Ctrl*: Unc13A and -B null animals expressing wildtype Unc13A; *CaM*$^{W1620R,W1622R}$ mutant: Unc13A and -B null animals expressing the Unc13A *CaM*$^{WRWR}$ mutant. Number of NMJs/cells (n) and animals (N): *Ctrl*: n=13, N=5; *Unc13CaM*$^{WRWR}$ *mutant*: n=14, N=5. Scale bar indicates 5 µm.

**Fig. S3**

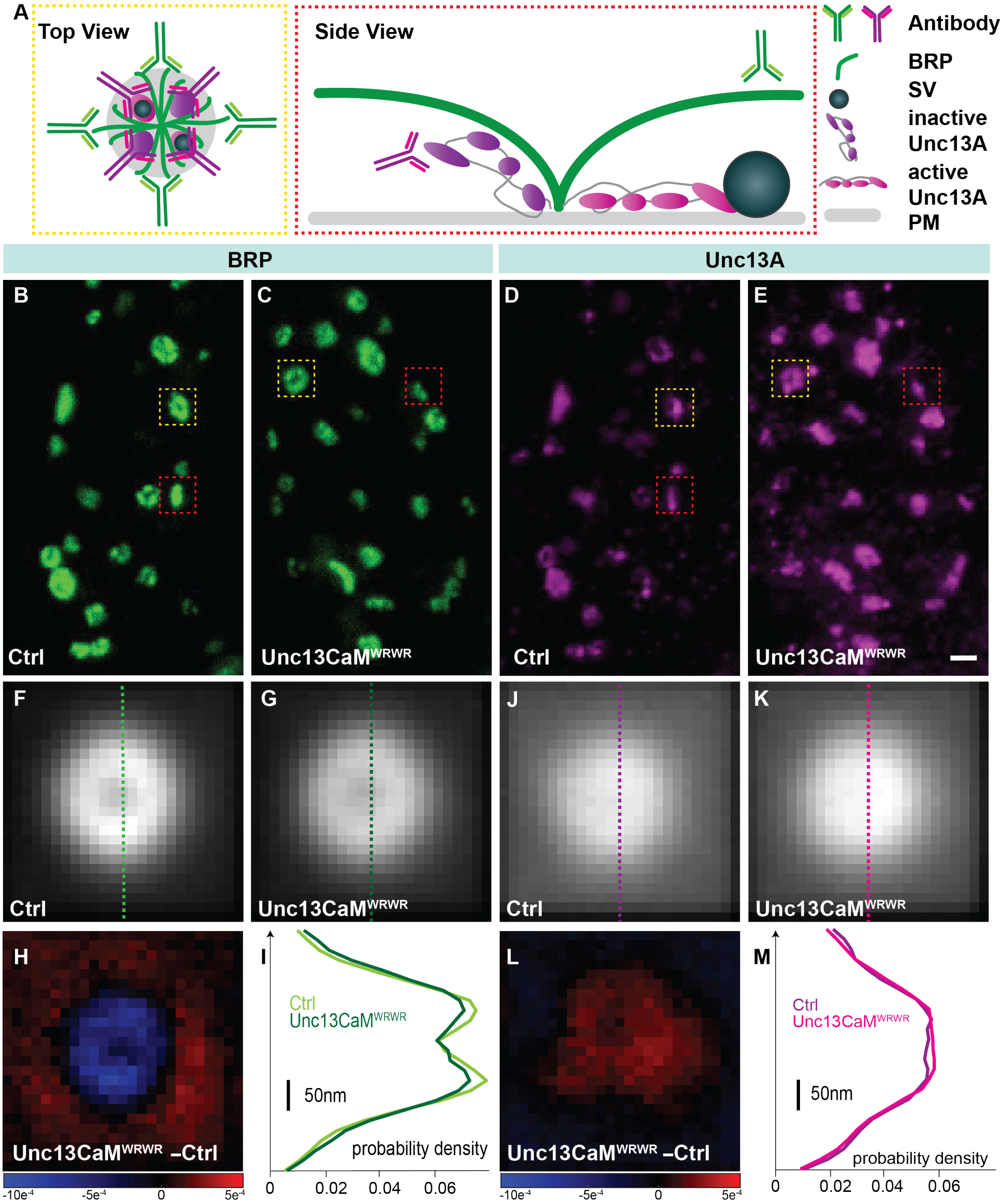


**Fig. S3 Top viewed Unc13A shows no changes in the position of the Unc13A C-terminus in the CaM mutant. (A)** Schematics of top (yellow dashed box) and laterally (red dashed box) viewed AZ with BRP in green and Unc13A in purple (inactive) or pink (active). Schematic antibodies indicating binding epitopes (green: NC82/Bruchpilot; purple: MUN-Domain). **(B-E)** STED images of muscle 4 NMJs of segment A2–4 from third-instar larvae of the displayed genotypes labeled with antibodies recognizing BRP (green) and Unc13A (purple). Red dashed boxes show laterally viewed AZ. Yellow dashed boxes show top viewed AZ. **(F-G)** Average and normalized BRP signal of the displayed genotypes. Both genotypes are displayed on the same scale. Dashed vertical lines through the midpoint of the image show which intensity values were used to generate line profiles (I). **(H)** Subtracted BRP signal displayed at the same scale as in Fig2H and L. **(J-K)** Average and normalized Unc13 signal of the displayed genotypes. Both genotypes are displayed at the same scale. Dashed vertical lines through the midpoint of the image show which intensity values were used to generate line profiles (M). **(L)** Subtracted Unc13A signal displayed at the same scale as in Fig2H and L. **(I-M)** Line profiles of BRP and Unc13A for the displayed genotypes.

Supplementary material

Profiles are scaled such that area under the curve equals 1. Genotypes (see methods for details): *Ctrl*: Unc13A and -B null animals expressing wildtype Unc13A; $CaM^{W1620R,W1622R}$ mutant: Unc13A and -B null animals expressing the Unc13A $CaM^{WRWR}$ mutant. Number of AZs ($\nu$), NMJs/cells (n) and animals (N): *Ctrl:* $\nu$ = 229, n=9, N=4; $CaM^{WRWR}$ mutant: $\nu$ = 224, n=7, N=4. Scale bar: 400 nm.

Supplementary material

**Fig.S4**

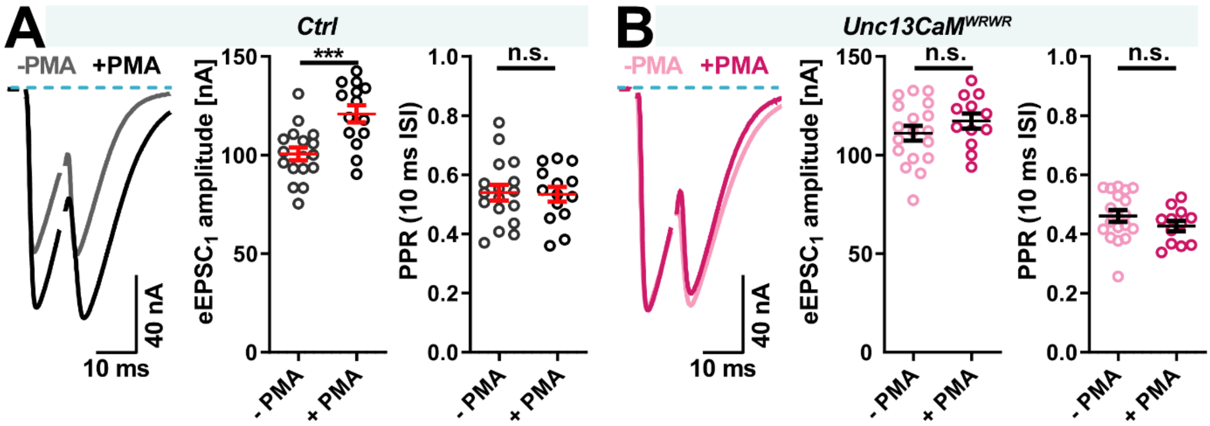


**Fig. S4: Respective occlusion of presynaptic potentiation by activation of the Unc13A CaM and C1 domains.** Panels (**A - B**) show data obtained in TEVC experiments from larval M6 NMJs at 1.5 mM extracellular $Ca^{2+}$. Graphs indicate mean values ± s.e.m. with * p ≤ 0.05; ** p ≤ 0.01. (**A & B left panel**) Example traces of AP evoked paired-pulse responses preincubated with DMSO (- PMA) or PMA (+ PMA) in *Ctrl* (-PMA: 17 NMJs from 10 animals, + PMA: 14 NMJs from 9 animals) and *Unc13CaM$^{WRWR}$* (-PMA: 18 NMJs from 11 animals, + PMA: 12 NMJs from 7 animals). (**A & B, middle panel**) Quantification of $eEPSC_1$ amplitudes showing a significant increase of transmission in PMA treated *Ctrl* cells (**A, middle panel**) and no effect of PMA treatment in Unc13*CaM$^{WRWR}$* mutants. (**A & B, right panel**) Quantification of paired-pulse ratios (PPR, 10 ms inter-stimulus interval (ISI)) revealed no change in PPR values in *Ctrl* (**A, right panel**) and *Unc13CaM$^{WRWR}$* (**B, right panel**) NMJs upon PMA treatment. Genotypes (see methods for details): *Ctrl*: Unc13A and -B null animals expressing wildtype Unc13A; *CaM$^{W1620R,W1622R}$* mutant: Unc13A and -B null animals expressing the Unc13A *CaM*$^{WRWR}$ mutant.
